# Resolving somatic SVs via full-length sequence model-based local graph-genome optimization

**DOI:** 10.1101/2025.02.11.636543

**Authors:** Kailing Tu, Qilin Zhang, Yang Li, Yucong Li, Lanfang Yuan, Jing Wang, Jie Tang, Lin Xia, Wei Huang, Dan Xie

**Affiliations:** Laboratory of Omics Technology and Bioinformatics, Frontiers Science Center for Disease-related Molecular Network, State Key Laboratory of Biotherapy, West China Hospital, Sichuan University, Chengdu, Sichuan, 610041, China; School of Mathematics and Statistics, Key Laboratory for Applied Statistics of the Ministry of Education, Northeast Normal University, Changchun 130024, China

**Author notes:** Corresponding authors: Laboratory of Omics Technology and Bioinformatics, Frontiers Science Center for Disease-related Molecular Network, State Key Laboratory of Biotherapy, West China Hospital, Sichuan University, Chengdu, Sichuan, 610041, China. No. 17, Section 3, Renmin South Road, Chengdu, Sichuan 610041, China. (D.X.), School of Mathematics and Statistics, Key Laboratory for Applied Statistics of the Ministry of Education, Northeast Normal University, 5268 Renmin Street, Changchun, Jilin Province 130024, China. (W.H.).

## Abstract

Somatic structural variations (SVs) are critical genomic alterations in cancer genomes. Long-read sequencing (LRS) is theoretically optimal for detecting somatic SVs. However, influenced by reads-to-reference alignment errors, particularly in low-complexity or highly repetitive genomic intervals, current LRS-based somatic SV callers fail to accurately detect SVs. Moreover, the lack of comprehensive ground-truth datasets hinders accurate evaluation. Here, we introduce SVscope, a novel algorithm that fundamentally addresses these challenges by leveraging full-length sequence information from span-reads and integrating local graph-genome optimization with a random forest strategy. SVscope outperforms state-of-the-art methods on six paired long-read whole-genome sequencing (WGS) benchmark cell lines, achieving a maximum F1-score improvement of 16.7%. In simulated datasets, SVscope demonstrates superior performance in both somatic SV detection and read phasing tasks. Based on the findings from SVscope, we validated 47 somatic SVs, thereby significantly expanding the existing experimentally validated ground-truth somatic SV dataset by 69.1%.

## Introduction

Somatic structural variations (somatic SVs) or de-novo SVs can be broadly defined as large genomic alterations exceeding 50 bp in length that occur in specific somatic cell clones. Somatic SVs, which involve larger segments of base variations compared to point mutations, are predominantly derived from genomic rearrangement events ^1, 2^. Existing studies have categorized SV events into five basic types based on their impact range: large-scale chromosomal rearrangements or breakend (BND) SV events, including duplications (DUP), inversions (INV), and translocations (TRA); and small-scale local genomic sequence alterations, including insertions (INS) and deletions (DEL) ^3^. These SVs tend to have a greater effect on gene expression than point mutations^4, 5^. Recent studies have identified that somatic SV events mediate genomic copy number variations (CNVs)^6^, gene fusions^7^, and cis-regulatory alterations^8^. In addition to these mechanisms, somatic SV events also influence the topological structure of the genome by altering CTCF binding sites, thereby participating in long-range gene regulation^9^, and modulate the methylation status of genome^10^. Accurate detection of somatic SVs is of great importance for discovering potential therapeutic targets for tumors, elucidating the regulatory networks of tumor genomes, and understanding the evolutionary patterns of cancer.

The scope of somatic SV detection task is to identify variants unique to somatic genome. Typically the detection process includes reads-to-reference alignment, abnormal alignment breakpoint extraction and somatic event detection. Second-generation sequencing (SGS) techniques, constrained by the short length of sequence reads that often fail to span entire SV regions. SGS-based algorithms predominantly relied on inter-read information arising from read populations, including alignment breakpoint ^11-13^ and sequencing depth^14, 15^ to decipher somatic SVs. Long-read sequencing (LRS) data has revolutionized this field by providing full-length sequence that spans entire SVs thereby enabling the precise detection of somatic SVs that were previously undetectable by second-generation sequencing (SGS)^3^. Leveraging the extensive sequence information from each individual read, current LRS-based algorithms utilize the intra-read information for somatic SV detection. To enhance detection accuracy, these methods focus on improving the extraction of alignment breakpoint features, with innovations including breakpoint clustering-based somatic genome polishing ^16^, tandem repeat sequence breakpoints merging ^17^, and sequence-to-image approaches ^18^. Alignment breakpoint information is the key reflection of SV events in the alignment results. For large-scale chromosomal rearrangement SV events including TRA, INV, and DUP, long-read sequences are decomposed by aligners into several independent alignment records, manifesting as split-alignment signature^19, 20^. Since these split-alignment somatic SVs typically do not involve local sequence gain or loss, breakpoint coordinates information is powerful for yielding accurate results in these large-scale structural variation detection tasks.

In contrast, for small-scale somatic SVs, like local INS and DEL, breakpoint information primarily originates from large segments of insertion or deletion in the read alignment CIGAR values, manifesting as an inner-alignment signature. Relying solely on breakpoint position information faces two challenges in the inner-alignment somatic SV detection: incomplete information and ambiguous localization. On the one hand, inner-alignment somatic SVs generally involve local sequence loss or gain, where a single breakpoint position cannot fully describe all information of the SV event. On the other hand, influenced by local variable number tandem repeat (VNTR) sequences, reads supporting the same SV event may produce multiple breakpoint features^21^. In addition to the aforementioned issues, the misalignment of reads in regions with highly repetitive sequence, including alpha-satellite DNA from centromeres^22-24^ and conserved tandem repeat sequences from telomeres^25^, also serves as a source of error in somatic SV detection tasks. Furthermore, the current landscape is marked by a lack of large-scale ground-truth somatic SV datasets. Past evaluations of somatic SV callers have predominantly relied on benchmark datasets derived from multi-omics data and the consensus of somatic SV callers. However, limitations in methodological design can lead to the omission of certain somatic SV events, thereby affecting the accuracy and comprehensiveness of somatic SV assessments. Despite these challenges, long-read sequencing offers unique advantages that lies in its ability to capture the full-length sequence spanning the entire somatic SV event and its flanking regions. Therefore, a systematic approach combining alignment breakpoint and full-length sequence information is essential for precise somatic SV detection.

To address this need, we introduce SVscope, a computational framework that leverages full-length sequence information and local graph genome optimization to accurately detect somatic SVs. The framework utilizes read alignment breakpoint information from the whole-genome scale to cluster and identify split-alignment somatic SVs and candidate inner-alignment somatic SVs. To mitigate the impact of alignment errors on inner-alignment somatic SV detection, SVscope re-analyzes the alignment relationships among all full-length sequences spanning the candidate somatic SV interval using a partial order alignment (sPOA) graph with multi-sequence alignment representation^26^ and accurately clusters reads with a sequence mixture model. To avoid read coordination errors affected by centromeres, telomeres, and segmental duplication sequences, SVscope also implements a random forest machine learning approach based on local alignment features to filter high-confidence somatic SV events. Evaluated on Oxford Nanopore whole-genome sequencing datasets from six tumor-normal benchmark cell line pairs, as well as simulated datasets with high heterogeneity in tandem repeat regions, SVscope outperformed existing state-of-the-art somatic SV callers. Based on the result of SVscope, we validate 47 somatic SV events with PCR methods improving the volumns of exprimental validation dataset by 69.1%. Furthermore, SVscope provides a single-read resolution visualization tool, ScopeVIZ, to facilitate detailed analysis of somatic SVs. All code implementations are publicly available on GitHub (https://github.com/Goatofmountain/SVscope).

## Result

### Error analysis of somatic SV Detection in cell lines

To elucidate the challenges in somatic structural variant (SV) detection, we conducted an in-depth analysis of existing multiplatform whole-genome sequencing datasets from six tumor-normal paired immortalized B lymphocyte cell lines (Supplementary Table S1). Among these, only COLO829 has a gold-standard somatic SV dataset based on extensive experimental validation^27^. The remaining five tumor-normal cell line pairs lack such golden-standard datasets. Therefore, we first constructed a benchmark dataset of somatic SVs for these cell lines by identifying recurrent somatic SV events across 3 sequencing platforms and 8 somatic SV callers (Supplementary Figure S1A, Methods). Our analysis revealed distinct patterns: HCC1395, H2009, and HCC1937 predominantly exhibited INS and DEL, which are categorized as inner-alignment somatic SVs, while HCC1954 and H1437 showed a higher prevalence of DUP, INV, and TRA, which are classified as split-alignment somatic SVs (Supplementary Figure S1B).

Four out of five existing somatic SV callers exhibit higher detection accuracy for split-alignment somatic SVs compared to inner-alignment somatic SVs (Figure 1A, Supplementary Figure S2). To further investigate the challenges in somatic SV detection, we analyzed the recurrent false positives (FP) and false negatives (FN) across different cell lines using five LRS-based somatic SV callers. Our findings indicated that low-complexity and tandem repeat sequences are hotspots for both FP and FN errors, with an enrichment of inner-alignment SV errors in these intervals(Figure 1B). This enrichment is likely due to the influence of the minimizer-and-chain algorithm used by long-read aligners, where local loss or gain of low-complexity or tandem repeat sequences can result in multiple breakpoint locations, leading to significantly larger deviations between the breakpoint coordinates and the true SV center positions compared to split-alignment SVs (DEL: p-vlaue=3.9e-6, INS: p-value=5.5e-5,Mann-Whitney U Test, Figure 1C). In contrast, split-alignment SVs, which rarely involve local sequence loss or gain, show minimal deviation in breakpoint coordinates in such regions (Figure 1C). This observation likely explains why most of breakpoint-based methods achieve higher precision in detecting these types of SVs.

**Figure 1.**
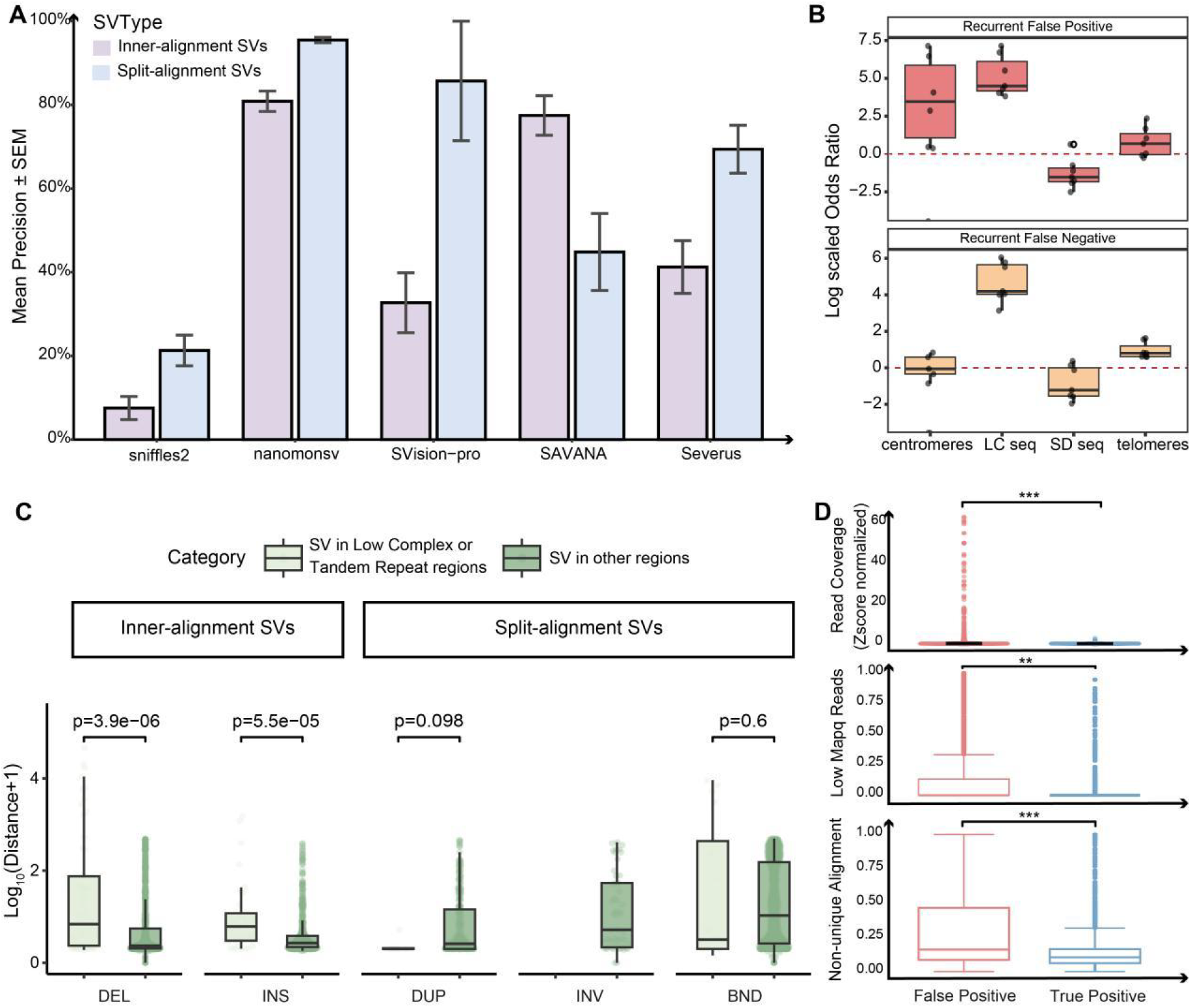
Error summary of somatic SV calling task. A. Barplot shows the mean precision of 5 existing somatic SV callers in detecting inner-alignment SVs (light purple) and split-alignment SVs (light blue) tasks. B. Boxplot shows the log scaled odds ratio of recurrent false positive (light red) and recurrent false negative (light yellow) somatic SVs in four types of reference genome. These regions include centromeres (indicating the centromeres regions of chromosomes), telomeres (representing the telomeric regions of chromosomes), SD seq (denoting the segmental duplication regions), and LC seq (referring to the low-complexity and tandem repeat sequence regions annotated by RepeatMasker software in the human reference genome). C. Boxplot shows log scaled breakpoint deviation of somatic SV located within low-complexity or tandem regions (light blue) and other regions (dark blue). D. Boxplots show the difference of local read coverage (upper), percentage of low mapping quality read (middle), and rate of non-unique alignment reads (lower) in false positive (light red) and true positive (light blue) somatic SV intervals with statistical significance (t-test) annotated as ns (p > 0.05), * (p < 0.05), ** (p < 0.005), and *** (p < 0.0005).

Beyond low-complexity and tandem repeat sequences, we also observed an enrichment of false positive somatic SV events in centromeres and telomeres (Figure 1B), which are highly repetitive regions composed of large segments of densely arranged, highly homologous tandem repeat sequences. These regions, characterized by their repetitive nature, pose significant challenges for accurate SV detection. For instance, centromeres are primarily composed of alpha-satellite DNA, which consists of arrays of ∼171 base-pair units arranged into higher-order arrays throughout the centromere of each chromosome^22-24^. Similarly, telomeres contain conserved tandem repeat sequences that are critical for maintaining chromosome stability^28^. The repetitive nature of these sequences leads to alignment ambiguities and errors. Reads that fail to span these regions would non-uniquely align multiple times to different genomic intervals with similar sequences, resulting in high local coverage depth and a high rate of low mapping quality reads (Supplementary Figure 3A-B). As a result, compared to true positive somatic SVs, recurrent false positive SVs in these regions exhibited higher local read counts (p-value < 2.2e-16, Mann-Whitney U Test Figure 1D, Supplementary Figure 3C), a higher proportion of low mapQ reads (p-value <2.2e-16, Mann-Whitney U Test, Figure 1D, Supplementary Figure 3D), and a higher proportion of non-uniquely aligned reads (p-value <2.2e-16, Mann-Whitney U Test, Figure 1D, Supplementary Figure 2E). These characteristics reflect the alignment inaccuracies caused by long-read sequences that cannot effectively span these regions, thereby significantly impacting the accuracy of somatic SV detection. In summary, our analysis highlights the complexities and challenges in somatic SV detection, particularly in regions with low-complexity, tandem repeats, centromeres, and telomeres. The limitations of current SV callers in accurately detecting inner-alignment SVs in these regions underscore the need for a more robust and accurate detection approach.

**Figure 2.**
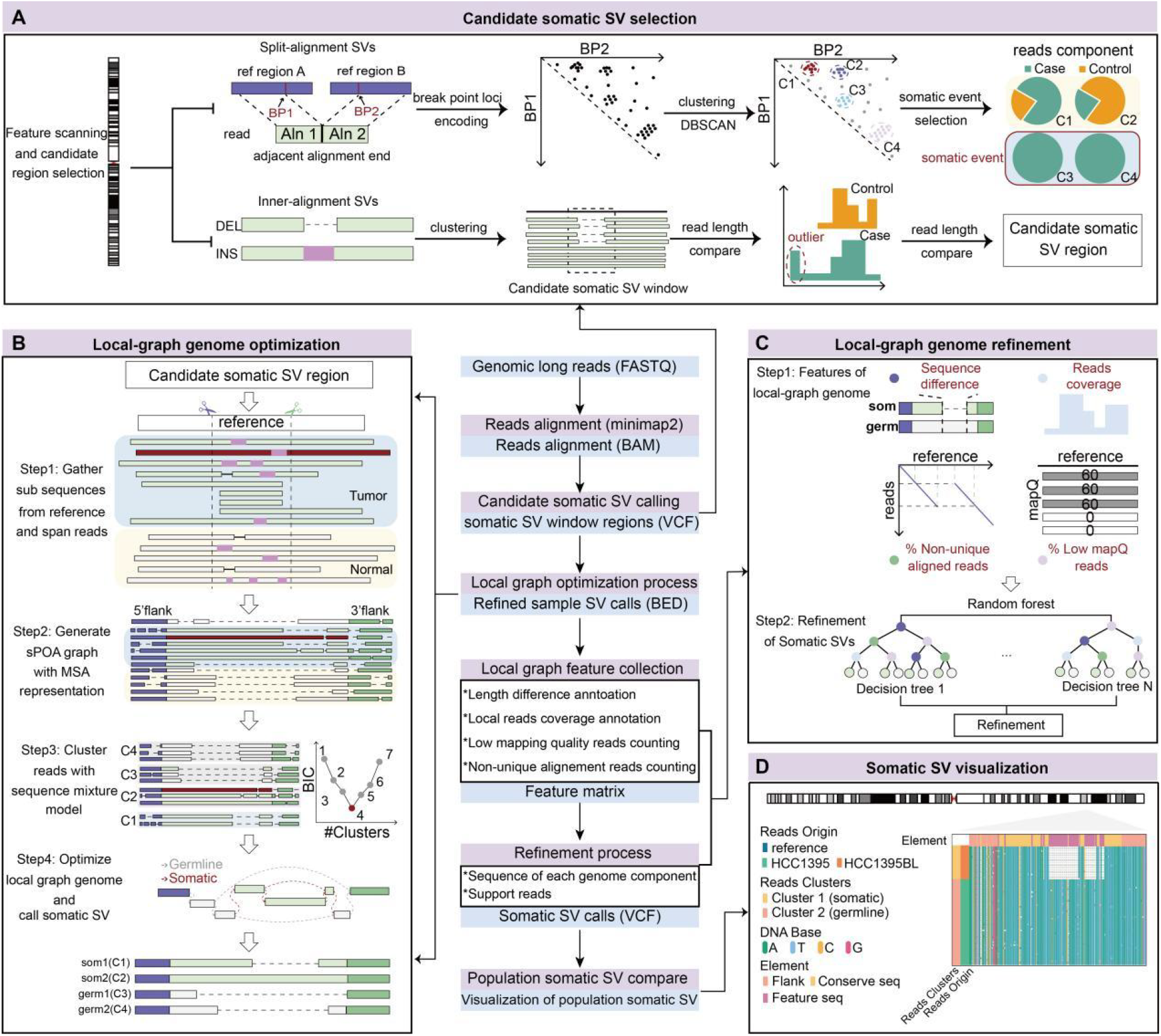
Schematic of the SVscope Workflow. This schematic illustrates the workflow of the SVscope algorithm for detecting somatic structural variations (SVs) from FASTQ data and comparing somatic SV sequences across population samples. The detection process consists four modules: A. Candidate somatic SV selection module selects candidate somatic SV intervals through paired-breakpoint clustering (split-alignment SVs) and full-length sequence length comparison (inner-alignment SVs). B. Local-graph genome optimization utilizes the full-length sequence information of span-reads to optimize local graph-genome and find case-reads-specific paths as somatic SVs. C. Local-graph genome refinement module evaluate the confidence of local-graph genome with a random forest model based on local alignment features. D. Somatic SV visualization module, visualize read contributions in SV detection.

**Figure 3.**
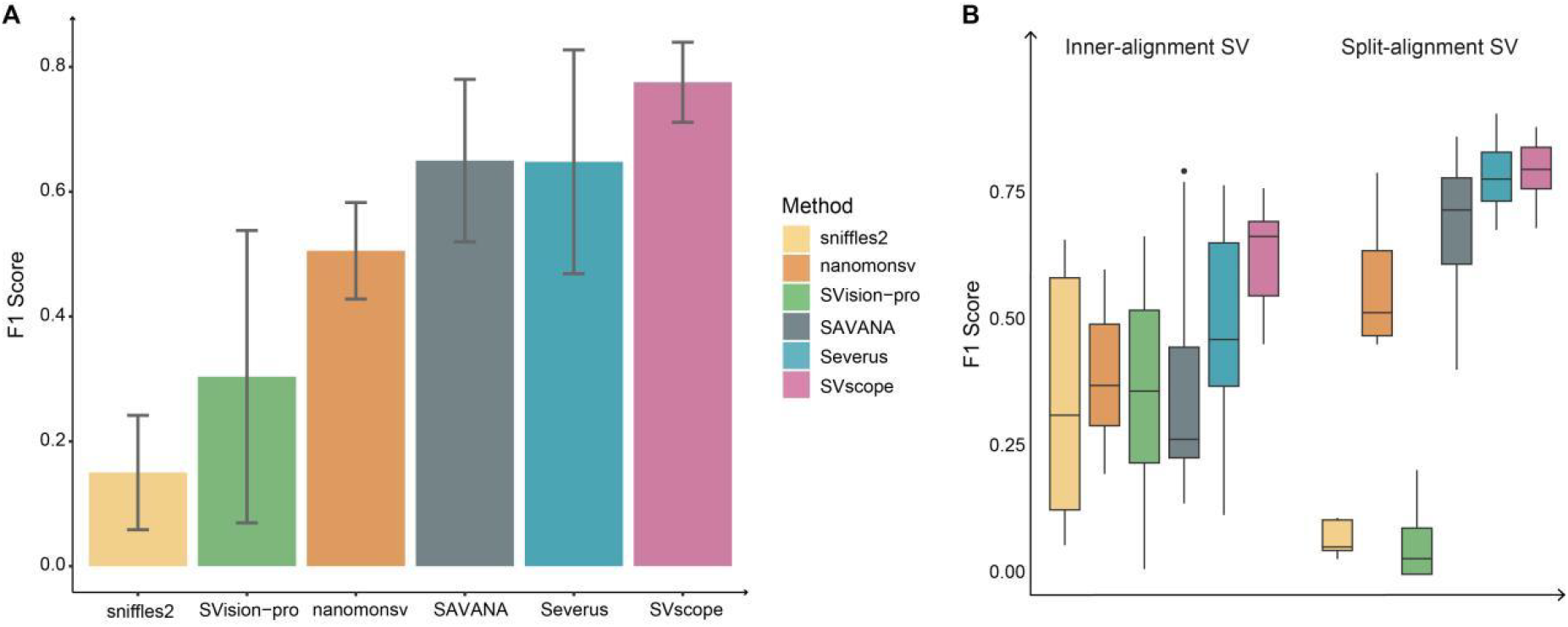
Performance of somatic SV caller in benchmark cell lines. A. Bar plot showing the F1 scores for different cell lines, comparing the performance of SVscope with other callers. B. Box plot displaying the percentage of somatic SVs detected near low-complexity sequences by various callers. The box represents the interquartile range (IQR), with the line inside the box showing the median. Whiskers extend to the most extreme data points not considered outliers, and outliers are plotted as individual points.

## Design of SVscope

To address the challenge of accurate somatic SV detection, refinement, and visualization, we introduce SVscope, which consists of four essential modules. First, the candidate somatic SV selection module is designed to analyze read alignment results and identify candidate somatic SV intervals. It categorizes potential somatic SVs into split-alignment SVs and inner-alignment SVs based on breakpoint characteristics (Figure 2A). Split-alignment SVs are characterized by reads with different segments aligning to multiple positions in the reference genome, and identified through paired-breakpoint coordinate, re-encoding, clustering and component analysis process (Methods, Supplementary Figure S4A). Inner-alignment SVs, featuring internal breakpoints with sequence insertions or deletions, are processed by clustering adjacent breakpoints to form candidate regions. The module then compares the lengths of case and control reads within these regions using full-length information from long reads to identify candidate somatic SV intervals (Methods, Supplementary Fig S4B). Second, the local-graph genome optimization module targets all inner-alignment somatic SV intervals. It constructs a local graph genome based on the complete sequences of span-reads from case and control samples using sPOA algorithm^26^, and optimizes it with a sequence mixture model (Figure 2B, Methods). Candidate somatic SVs are selected from case-read-specific paths in the local graph genome. Third, to mitigate read misalignment issues caused by highly repetitive regions, the local-graph genome refinement module employs a random forest model in conjunction with read alignment features to assess the accuracy of all previously obtained local-graph genomes, thereby filtering out high-confidence somatic SVs (Methods, Supplementary Figure S5A-B). Finally, to facilitate a clear comparison of read contributions in SV detection process, SVscope includes a somatic SV visualization module that enables users to inspect the clustering of all span reads (Figure 2D).

## Accurate resolve somatic SVs with SVscope

To evaluate the performance of SVscope in somatic SV detection, we sourced 7 pairs of tumor-normal cell ONT whole-genome sequencing datasets from public databases (Supplementary Table S1). SVscope identified five basic types of somatic SVs and showed superior overall performance compared to other somatic SV callers across all cell lines (Figure 3A, Supplementary Figure S6A), achieving a maximum F1 score improvement of 17.01% over the second-ranked somatic SV caller (Figure 3A, Supplementary Figures S6B-C). Additionally, SVscope exhibited robust detection efficiency in both inner-alignment and split-alignment SV tasks (Figure 2B, Supplementary Figure S6D).

To further assess SVscope’s detection capability for somatic SVs in low-complexity and tandem repeat regions, we constructed a simulated dataset centered around tandem repeat sequences in the human genome (Supplementary Figure S7, Methods). Under varying tumor purity and SV length conditions, SVscope consistently outperformed other somatic SV callers (Figure 4A, Supplementary Figures S8A-C) and maintained high-precision results across different tumor purity levels (Figure 4B). Notably, this precision advantage persists even when facing the added complexity of diploid haplotype interference, a critical challenge in distinguishing somatic variants from germline structural polymorphisms in repetitive regions. This demonstrates SVscope’s robustness against the influence of multiple-allele haplotypes compared to other alignment-based somatic SV callers (Supplementary Figure S8D). Beyond somatic SV event identification, SVscope also exhibits superior performance in the calculation of somatic SV support reads (Figure 4C) and provided robust estimations across different tumor purities (Supplementary Figure S9).

**Fig 4.**
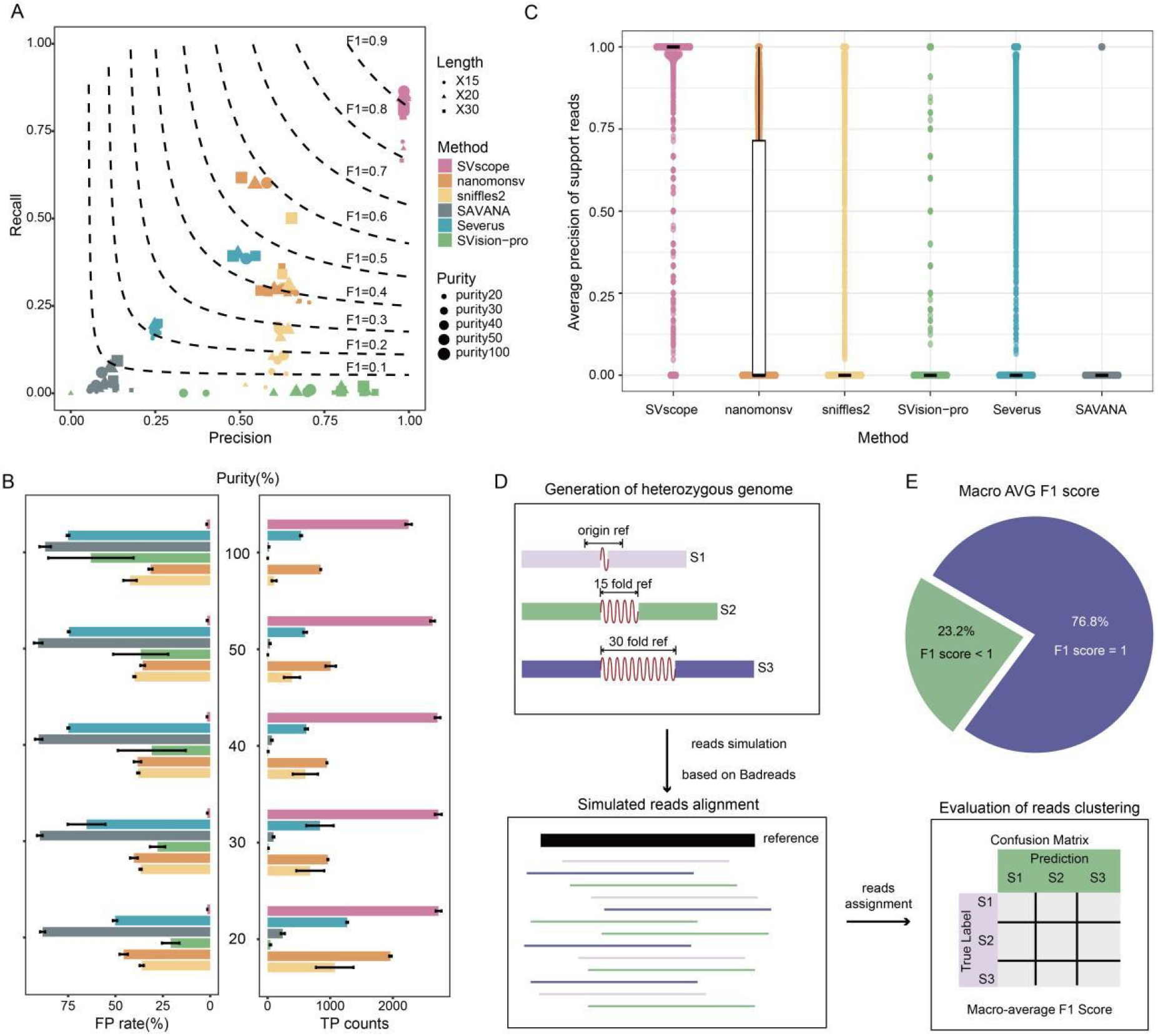
Performance of somatic SV callers in simulated somatic tandem repeat dataset. A. Scatter plot with contour lines illustrating the precision and recall of SVscope and other state-of-the-art somatic SV callers across all simulated dataset with 5 tumor purity threshold and 3 tandem repeat extension scale. The parameters of datasets and somatic SV callers are denoted on the right. Dashed lines represent various gradients of F1 scores. B. Barplot shows the count of true positive somatic SVs and false positive rate of each somatic SV callers under different tumor purity. C. Scatter plot shows the accuracy of somatic SV support reads in each somatic SV callers under different tumor purity. tumor purity and SVcaller name in Fig.4B-C are color-coded in Fig.4A. D. Schematic of simulated data for heterogenous genome. E. Pie chart shows the percentage of simulated heterogenous regions with perfect read assignment performance using SVscope (F1 score=1).

The employment of full-length sequence based local-graph optimization strategie is critical in somatic SV support read detection tasks. For instance, a long tandem repeat somatic insertion interval on chr22:50137660-50137991 illustrates how reads harboring somatic SVs are prone to alignment breakpoint errors due to the variable choice of minimizers in the nearby flanking sequences, leading to different alignment breakpoints for reads originating from the same somatic genome (Supplementary Figure S10A). In addition to the high-dispersion INS breakpoints, such long tandem repeat insertions can even introduce duplication-like breakpoints primarily manifested as supplementary alignments (Supplementary Figure S10B). Thanks to the application of full-length span-read sequences and sPOA graph-based multiple sequence alignment strategies, SVscope is unaffected by such read-reference alignment errors, ensuring proper read alignment (Supplementary Figure S10C). Based on this, the local graph genome optimization process employing the sequence mixture model can accurately estimate the genomic information of different cellular components, thereby enabling precise parsing of heterogeneous genomes (Supplementary Figure S10D).

To further test SVscope’s ability to distinguish read origins in heterogeneous genomes, we mixed two simulated somatic genomes with different tandem repeat lengths (S2: X15, S3: X30) and a control genome (S1). We then used SVscope to perform phasing calculations on the reads and evaluated the accuracy of the phasing results (Figure 4D). In this task, SVscope achieved completely accurate phasing results in 2359 out of 3225 (76.8%) simulated mixed heterogeneous genomes (Figure 4E). Taking the heterogeneous simulated region chr22:10,538,368-10,538,453 as an example(Supplementary Figure S11A), SVscope effectively mitigates the impact of local breakpoint errors, accurately clustering reads through its full-length sequence (Supplementary Figure S11B) and local graph genome optimization process (Supplementary Figure S11C-D). This performance underscores SVscope’s robustness in accurately clustering reads from different genomic origins, highlighting its potential for reads phasing tasks.

## Application of SVscope in somatic SV detection and validation

Given SVscope’s superior performance in somatic SV detection tasks across both cell lines and simulated datasets, we further optimized the gold-standard somatic SV dataset for the HCC1395 cell line. Based on SVscope, we identified 262 somatic SV events not present in the existing benchmark datasets from the HCC1395 cell line, comprising 149 inner-alignment SVs and 113 split-alignment SVs (Figure 5A). Among these, 114 out of 262 somatic SV events were uniquely detected by SVscope (Supplementary Figure S12). These SVs exhibit complex breakpoint distributions. For example, a somatic tandem repeat insertion event within the intron of the CNGA2 gene located at chrX:151,742,802-151,743,170 manifests multiple alignment breakpoints in the read alignments of HCC1395, with longer insertion fragments ranging from 62 to 81 bp and shorter fragments ranging from 8 to 34 bp (Figure 5B). These short fragment breakpoints are often overlooked by other somatic SV callers. In contrast, HCC1395BL shows a single alignment breakpoint with insertion fragment lengths ranging from 56 to 60 bp (Figure 5B). Visualization results from ScopeVIZ indicate that reads originating from HCC1395 are longer than those from HCC1395BL (Figure 5C). Through the local graph genome optimization process, we reconstructed the genomic sequences of HCC1395 and HCC1395BL in this region (Figure 5D). PCR experimental gel electrophoresis also reflects bands consistent with SVscope’s computational results (Figure 5E). Similarly, in the case of a somatic deletion SV within the intron of the EXOC2 gene located at chr6:599,529-604,175, SVscope demonstrates robust detection and read clustering capabilities (Supplementary Figure S13). The breakpoints of these SVs exhibit highly dispersed distributions in the read alignments, and the paired HCC1395BL cell line also harbors similar sequence insertions or deletions relative to the reference genome. These features interfere with other alignment-based algorithms, causing them to fail to accurately identify such SVs. In total, based on SVscope’s computational results, we validated 47 somatic SV events, 34 of which were not present in the benchmark datasets with 10 of them uniquely detected by SVscope (Supplementary Figure S14, Supplementary Table S2, Extend Data File 1). These validated events significantly enhance the existing somatic SV dataset for the HCC1395 cell line, providing a robust foundation for future research.

**Figure 5.**
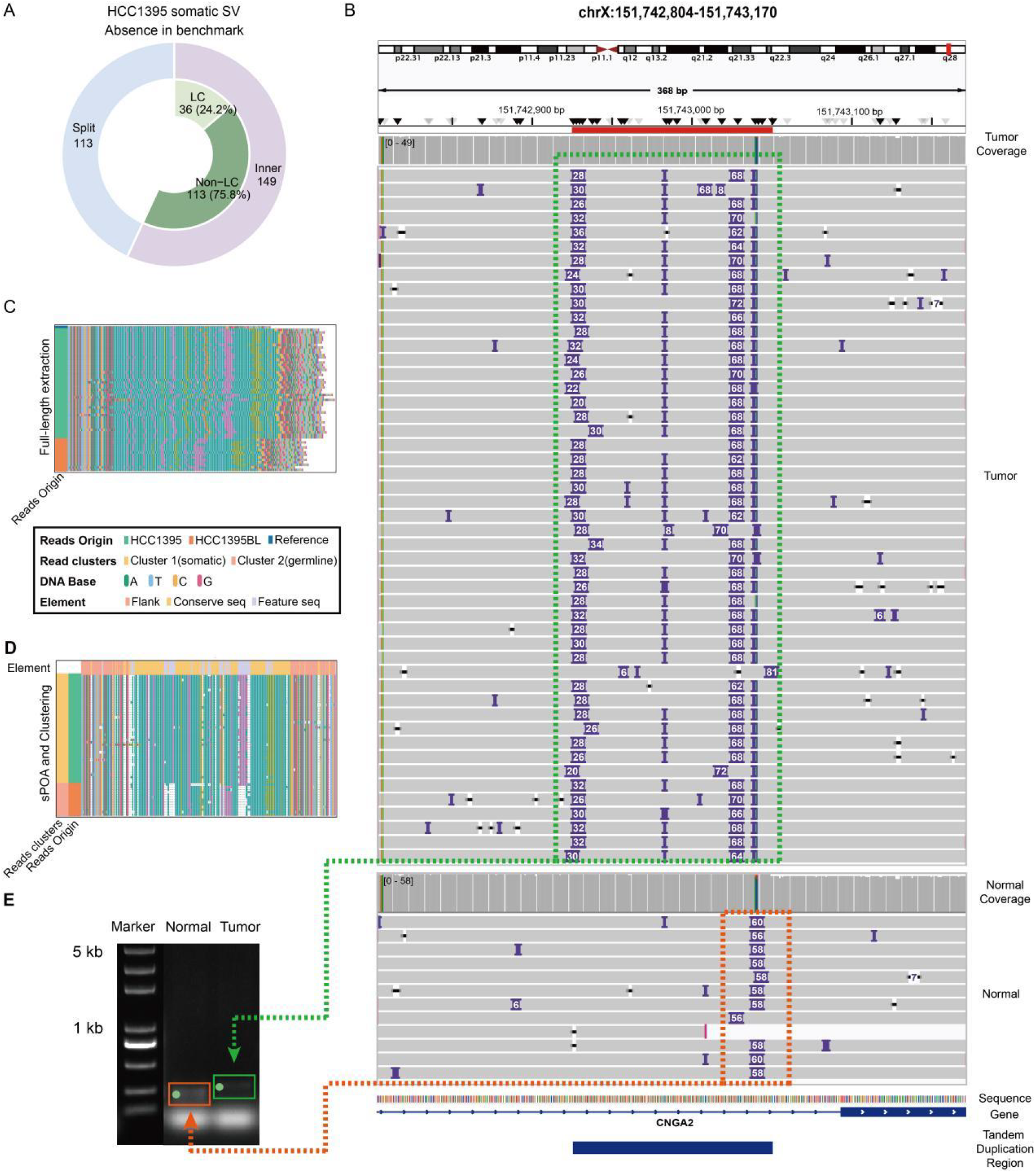
Detailed characterization and validation of somatic SV events. A. The outer pie chart shows the percentage of inner-alignment (purple) and split-alignment (light blue) somatic SVs identified by SVscope from HCC1395 cell line that is abcense in benchmark SV sets. The inner pie chart shows the percentage of low-complexity or tandem repeat sequence associated (light green) and non-associated (dark green) somatic SVs events. B. Genome browser plot shows the example somatic SV region chr6:600,931-602,770 in HCC1395 (top) and HCC1395BL (bottom). C. Bar plot shows the raw sequence extracted from HCC1395 (colored in green) and HCC1395BL reads (colored in orange). D. Bar plot shows the result of reads clustering based on SVscope. E. The gel electrophoresis image illustrates the targeted PCR results for the target somatic SV events in the genomes of HCC1395BL (left panel, Normal) and HCC1395 (right panel, Tumor). The target bands are highlighted with green dots.

## Discussion

SVscope introduces a novel computational approach that combines local graph genome optimization and random forest machine learning strategies, representing a significant advancement in the field of somatic structural variant (SV) detection. Specifically, SVscope leverages full-length sequence information from span reads and integrates it with a local graph genome optimization process. This approach effectively mitigates alignment errors introduced by read aligners, thereby ensuring the accuracy of the algorithm. Additionally, the employment of a machine learning strategy based on local alignment features reduces false positives caused by alignment errors in highly repetitive regions, such as centromeres and telomeres. These innovations collectively enhance the robustness and precision of somatic SV detection and read phasing tasks. SVscope provides a more accurate and reliable tool for analyzing complex and heterogeneous genomes, paving the way for future advancements in genomic research and clinical applications.

The development of SVscope highlights the potential of leveraging full-length information to improve the accuracy and reliability of somatic genome analysis. This approach not only enhances the detection of somatic SVs but also provides valuable insights into the heterogeneity of complex genomes. Beyond the nucleotide sequence, the integration of extensive epigenetic data including methylation^29,30^ and chromatin accessibility^31-34^ from single-molecule offers a more comprehensive view of the genome, which is crucial for understanding the underlying mechanisms of genetic disease and cancer. Future research should focus on optimizing algorithms to further explore the potential functions of somatic SVs, leveraging the multi-dimensional information provided by long-read single-molecule sequencing technologies.

The local graph genome optimization step is the core component of the SVscope algorithm, relying on span-reads to provide full-length sequence for the target genomic intervals. However, due to the limitations of read length in current long-read sequencing technology, somatic SVs with ultra-long insertions cannot always be precisely analyzed through SVscope due to the unavailability of span-reads. Advances in ultra-long read sequencing technologies^35^ are expected to mitigate these challenges in the future. The computational resource consumption is another significant issue with the current algorithm. This is primarily attributed to the sPOA-graph algorithm and the extensive sequence feature extraction required for the Expectation-Maximization (EM) algorithm in the sequence mixture model. Future research needs to focus on simplifying sequence feature extraction strategies. DNA large language models, including Evo^36^, Nucleotide Transformer^37^, and DNABERT^38^, have been proven to extract DNA sequence features and represent a promising direction for development. However, the lack of training sets currently limits the application of large language models for somatic SV detection tasks. The precise read clustering information provided by SVscope could serve as an important source of training sets for such large language models.

In summary, SVscope has demonstrated superior performance in somatic SV detection, SV support reads calculation, and reads phasing tasks. It has identified new somatic SV events in paired tumor-normal cell lines, which were validated through PCR experiments, thereby contributing to the construction of the gold-standard somatic SV dataset for the HCC1395-HCC1395BL benchmark cell line pair. Furthermore, the development of the ScopeVIZ visualization system allows users to visually inspect the contribution of sequencing reads to somatic SV detection, facilitating detailed confirmation of each event. SVscope thus provides a new paradigm for leveraging single-molecule long-read information, offering enhanced accuracy and interpretability in the analysis of somatic SV events.

## Methods

### Establishment of somatic SV benchmark dataset for cell lines via multiomics detection voting strategy

To create a benchmark dataset of somatic SVs for tumor-normal B lymphocyte pairs HCC1395-HCC1395BL, HCC1937-HCC1937BL, HCC1954-HCC1954BL, H1437-BL1437, and H2009-BL2009, we gathered high-depth whole-genome sequencing datasets obtained from Illumina short-read sequencing platforms and long-read sequencing platforms, including PacBio and Oxford Nanopore from published database. For short-read sequencing data, we employed three SV callers—SvABA, GRIPSS, and Manta—to detect somatic SVs. For long-read sequencing data, five SV callers—Sniffles2, SAVANA, Severus, SVision-pro, and NanomonSV—were used for somatic SV detection. The detection results were integrated using the Minda Ensemble module, and somatic SVs supported by at least four SV callers and data from at least two sequencing platforms were selected. For the COLO829-COLO829BL cell line pair, we directly adopted the somatic SV list provided in a previous study, which was validated through extensive experimental evidence, as the gold standard dataset. Large-scale somatic SV events including INV,DUP,and BND are labeled as split-alignment somatic SVs. Small-scaled somatic SV events including INS and DEL are labeled as inner-alignment somatic SVs.

### Evaluating Cell Line structural variants detection

To benchmark the performance of SV detection tools, we downloaded SV calls and ONT reads from five tumor cell lines: HCC1954, H2009, HCC1937, H1437, and HCC1395. The high-confidence calls for insertions, deletions, and duplications published by previous study served as our ground truth ^39^. We aligned the ONT reads to hg38 using minimap2(v2.26-r1175) with the default setting for SV detection. The somatic calling tools Nanomonsv(v0.7.0), Savana(v1.2.1), and Severus(v0.1.2) all follow the recommended workflows and default parameters. Although Sniffles2(v2.0.7) was not designed for a paired tumor-normal analysis, we utilized the recommended sniffles2-merge process and its mosaic mode to subtract the normal sample SV calls from the tumor sample calls. For all tools, the minimum supporting read count was set to 3, and the minimum SV length was established at 50 bp. To examine the correctness of detected SVs, Minda(v0.0.1)^39^ was employed to calculate precision, recall and F1-score between the ground truth and the call set, which is agnostic to SV types and is better suited for analysis of somatic SVs represented as junctions.

### Generation of simulated data

To benchmark the performance of SV detection tools, we simulated ONT sequencing data with a depth of 40× based on the human reference genome hg38 on chromosome 22. We designed five allele frequencies levels: 0.20, 0.30, 0.40,0.50and 1. Additionally, we included a control group without insertion variants and three new sequence insertion variants of different lengths: TD sequence X15, X20 and X30. The aim was to evaluate the accuracy, recall, and F1 scores of TD detection by Somatic SV callers under varying tumor purities, different new sequence insertion lengths, and the presence or absence of germline mutations. From the human repeat segment annotation database RepeatMasker, we retained TDs on chromosome 22 that are 1000 bp or shorter and ensured that there are no other TDs within 200 bp of each TD. This filtering resulted in a total of 6,478 candidate TD windows, with TD types including Simple_repeat and Satellite. These were divided into four files: A contains TD windows without any variations; B contains TD windows with germline variations; C contains TD windows with somatic variations; and D contains both somatic and germline variations. The positions and lengths of TDs in files C and D were recorded as a gold standard set for subsequent evaluations. First, using the simulation sequencing software VISOR (v1.1.2), we took chromosome 22 of the hg38 genome as the reference genome and used the recorded mutation files as input to generate corresponding variant genomes for different mutation types. Next, we employed the simulation sequencing software badread (v0.4.0) ^40^ (https://github.com/rrwick/Badread) to simulate sequencing at a depth of 40× for each mutation type, using the variant genomes generated by VISOR^41^ as the reference. This process produced a dataset of ONT simulated sequencing fragments with three lengths of new sequence insertions across five different tumor purity levels. The parameters for the badread simulation software were as follows: -quantity 40x --glitches 1000,20,20 -junk_reads 0.1 -random_reads 0.1 --chimeras 0.1 --identity 80,90,6 --length 20000,5000. The error model used was nanopore2023.

### Evaluating simulated structural variants detection

The ONT reads are aligned to the hg38 reference genome using minimap2 (v2.26-r1175) with default settings^42^. The somatic calling tools Nanomonsv, Savana, and Severus all follow the recommended workflows and default parameters. The minimum supporting read count was set to 3 for all tools, and the minimum SV length was set to 50 bp. Sniffles utilizes the recommended sniffles2-merge process from our previous study ^43^, while Straglr(v1.4.1) ^44^ adds the -Y parameter during alignment as required by the software, followed by the same merging operations as Sniffles. The merged BAM files are then used for Straglr with the following parameters: --genotype_in_size --min_support 3 --max_str_len 100 --max_num_clusters 9 --min_ins_size 50. The filtering process retains only the records supported by tumor reads. Tandem-genotypes(v1.9.0)^45^ runs according to the software’s default workflow and parameters. The results for case and control are analyzed using the Wilcoxon test and FDR correction, identifying records with at least 3 supporting reads and significant q-values as somatic.

To examine the correctness of detected somatic SVs, we employed region matching, which requires accurate detection of SV sites. For example, assume a particular benchmark somatic SV [b.chrom, b.start, b.end], and a prediction [p.chrom, p.start, p.end]; then a correct region-match should at least overlap 1bp. Any called somatic SVs without a matched prediction were counted as false negatives. Based on the numbers of true positives (TP) and false negatives (FN), we computed the recall, precision and F1-score with the following equations, respectively.

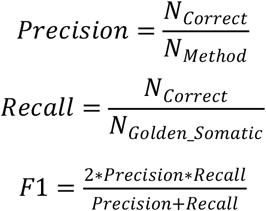

Each caller was run with different AF and insertion length of somatic SV, and the performance of detecting simulated SVs was assessed correspondingly.

### Validation of somatic SVs

DNA was extracted from HCC1395 and HCC1395BL using the QIAamp DNA Mini Kit (QIAGEN) according to the manufacturer’s instructions. The forward and reverse primers were designed according to the 2000bp upstream and downstream sequences of the aim somatic SV genomic intervals and synthesized by Sangon Biotech (Supplementary Table SXX Primers). Extracted DNA was subjected to PCR using matched primers. Each 50-μL reaction consisted of 2× Premix Taq (Takara) 25 μL, 2 μL 10 μM forward primer, 2 μL 10 μM reverse primer, 50-150 ng DNA, and add ddH2O to 50 μL. PCR was performed as follows: an initial step of 5 minutes at 95 °C, 30 cycles of 15 seconds at 95 °C, 15 seconds at 56 °C and 60-300 seconds at 72 °C (size of PCR products between 200 bp and 6 kb). Finally, the samples were incubated at 72 °C for 5 minutes. The PCR products were sent to Sangon for Sanger sequencing.

### SVscope methodology

#### Candidate somatic SV selection

The candidate somatic selection module of SVscope is the initial step in identifying candidate somatic SV loci. This process involves clustering alignment breakpoint information. To begin with, SVscope collects all read alignment information from both case and control samples. Subsequently, it extracts alignment breakpoint pairs from two sources: CIGAR values (for inner-alignment breakpoints) and supplementary alignment records (for split-alignment breakpoints). Each breakpoint pair i is encoded as below,

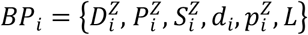

where status Z ∈ {0,1} represents the order of alignment breakpoint end with Z=0 for 5’ end, and Z=1 for 3’ end. The first 3 elements represents for the location of each breakpoint end in reference genome with D for contig ID, P for chromosome location, and S for the DNA strand. The last 3 elements represents for the location of each breakpoint end in read with d for readID, p for read location, and L for read resource.

For split-alignment somatic SVs, SVscope designs a systematic location encoder to further analyze their location relationships. Specifically, the chromosome IDs are sorted according to their IDs. Breakpoint positions on chromosomes with higher IDs are transformed into a unified coordinate system by summing the lengths of all preceding chromosomes. This approach ensures that breakpoint coordinates are consistently measured across different chromosomes, facilitating more accurate and systematic analysis of split-alignment somatic SVs. Paired split-alignment breakpoints which are closed in each read (by default located within 100bp in a single read) but different coordination in reference genome were re-encoded through this systematic location encoder and cluster through DBSCAN algorithm with default setting (distance=500 eps=3). Finally, only breakpoint clusters sourced from the case sample are identified as somatic SV events. This step ensures that the identified SV events are specific to the case sample and not present in the control sample, thereby improving the accuracy of somatic SV detection.

For deletion type inner-alignment breakpoints, SVscope merged all breakpoints from case reads to generate a candidate deletion set. For insertion type inner-alignment breakpoints, case read breakpoints located within Repeatmasker annotated tandem repeat regions were firstly merged into the target tandem repeat genome interval. The others would merged would be merged if the distance lower than preset threshold (by default 200bp). To select candidate inner-alignment somatic SVs with significant sequence loss or gain, all sub read sequence from both case and control samples spanning each candidate inner-alignment somatic SV were extracted with pysam packages. The candidate inner-alignment somatic SV intervals would be process into further analysis once case reads have larger (insertion case) or smaller (deletion case) length of span sequence over preset threshold (by default 50bp).

#### Local graph genome optimization

The candidate somatic SV selection module provides a set of candidate somatic SV genomic intervals. In the local graph genome optimization module, the SVscope algorithm targets all somatic structural variation (SV) candidate intervals and their flanking regions (by default capture a 50bp range upstream and downstream of the target somatic SV interval). Utilizing the pysam toolkit, it extracts sequence sets from the reference genome and reads spanning these intervals in both tumor and paired normal samples. For every sequencing read length within the candidate somatic SV interval, all alignment records are extracted based on their alignment coordinates. Alignments on the negative strand are converted to positive strand sequences using the pysam toolkit. Subsequently, SVscope constructs a local graph genome for each candidate somatic SV interval sequence set using the SIMID-sPOA algorithm and extracts the output results of its multiple sequence alignment representation. Highly heterogeneous alignment sites are filtered out to serve as features for subsequent multi-class sequence mixture models (second largest sequence frequency > 3 or frequency >= 5%). For the case of N reads and m highly heterogeneous alignment sites, we encoded data as an observed matrix,

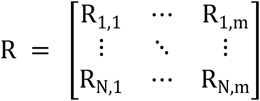

where one-hot vector 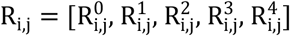 represents the observed value of the j-th feature in the i-th read, which is corresponding to the 4 DNA bases A,T,C,G and the alignment gap in the multisequence alignment representation of sPOA graph. The matrix encapsulates the presence or absence of specific nucleotide bases or gaps at each heterogeneous feature site across all reads.

#### Multi-category sequence mixture model

We consider each candidate somatic structural variation (SV) interval as a local graph genome assembly constituted by K types of lantent linear genomic sequences G = {G_1_, …, G_k_} . The composition of each genome in the whole cellular population is represented by the parameter π = [π_1_, …, π_k_] . Each linear genome G_r_ is represented as a set of m features, and all features constituting the local graph genome can be represented by the parameter matrix,

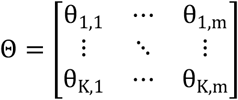

Where each element θ_i,j_ within Θ is composed of 5 probabilitie 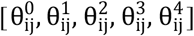, same as the observed data encode, corresponding to the four DNA bases A, T, C, G, and the alignment gaps.

For each sequencing read in the multisequence alignment representation of sPOA graph, R_r_ = [R_r,1,_ … , R_r,m_]we employ a multi-class sequence mixture model for modeling, denoted as,

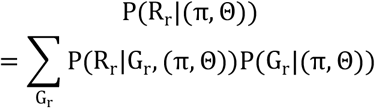

Then, the complete likelihood function of (R, G) is,

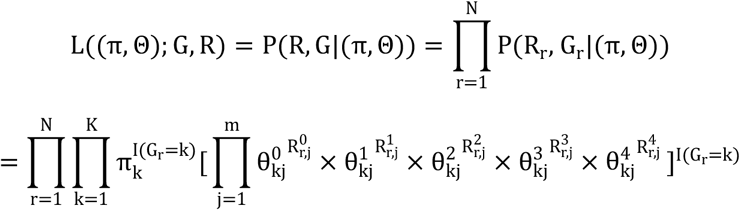

Therefore the log-likelihood function of the complete data is,

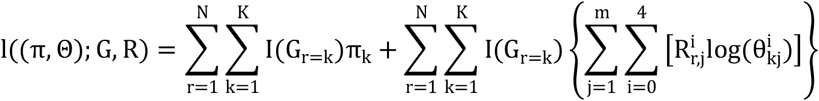

Parameter estimation. The SVscope algorithm initiates the parameter estimation process by employing hierarchical clustering to approximate the initial model parameters. Specially, SVscope leverages the scipy package (version 1.7.3) to perform hierarchical clustering on the reads with the edit distance matrix,

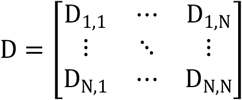

where 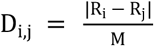, represents the edit distance between read i and read j. The outcome of this clustering, denoted as N × k one-hot matrix,

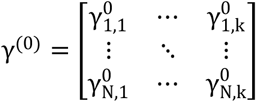

serves as the foundation for initial model parameters (π^(0)^, Θ^(0)^).

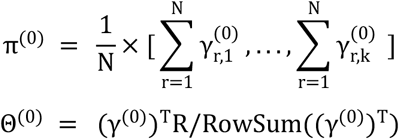

Next, SVscope algorithm commences the parameter estimation phase by employing the Expectation-Maximization (EM) algorithm to refine the parameters of the multi-category sequence mixture model (π, Θ). The Q function induced by the complete log-likelihood function is,

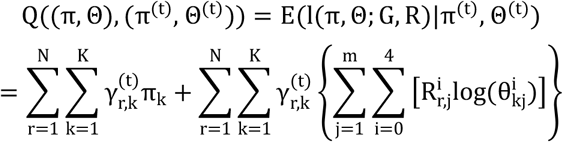

Where in the Expectation step,

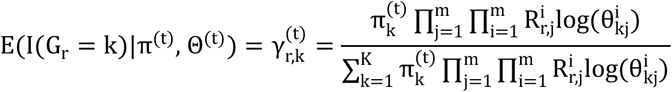

In the Maximization step,

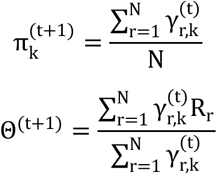

SVscope systematically estimates the model parameters for a spectrum of mixture model sizes, ranging from k=1 to k=10, and ascertain the optimal number of genome components K with minimal Bayesian information criterion (BIC),

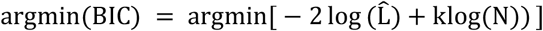

#### Local graph genome refinement

To evaluate the confidence of optimized local graph genome and avoid the false positive caused by highly-repeat region or segmental duplication sequence, we trained a Random Forest (RF) classifier using Scikit-Learn. To this aim, we selected 3,452 labeled candidate somatic SV intervals from five cell lines (HCC1395, H2009, H1437, HCC1954, HCC1937) using the local graph genome optimization module. These intervals were used as the candidate training set. We then summarized the local alignment features of these intervals as interval features. Specifically, we calculated the absolute sequence difference length between the tumor genome and the paired normal genome, marked as ABSMisScore. Additionally, we computed the proportion of reads spanning the interval in the case and control samples, the proportion of low mapping quality (<5) reads, the proportion of non-uniquely aligned reads, and the normalized coverage score based on the average and standard deviation of read counts in 10-kb windows across the genome. All data were randomly split into training and testing sets at a ratio of 4:1. The optimal values for these parameters were determined using cross-validation and grid search, with the number of decision trees ranging from 5 to 30 and the maximum depth ranging from 2 to 64. To detect somatic SVs in each cell line, we trained an RF model using the labeled SVs from all other cell lines to ensure that no data from the cell line to be analyzed was used during model training.

## Data and code availability

The sequencing data in this study is downloaded from SRA data base under accession number SRP494767 (BioProject: PRJNA1086849). Detail accession number for each cell lines were listed in Supplementary Table S1. Original code of SVscope and ScopVIZ are available from github at https://github.com/Goatofmountain/SVscope. Benchmark somatic SVs, including PCR validation label of HCC1395 cell line were listed in Extend Data File 1.

## Acknowledgments

This work was supported by grants from the National Natural Science Foundation of China (82173383 to D. X., 11671073 to W.H., 32300522 to K.T., 32200508 to L. X., 82072582 to J.T.), the 1·3·5 project for disciplines of excellence, West China Hospital, Sichuan University (ZYYC23024 to D. X.). This work is also supported by the Science and Technology Department of Sichuan Province, China (2025ZNSFSC1098 to K.T., 2024NSFSC1187 to Y.L.), and the Fundamental Research Funds for the Central Universities, China to W.H.

## Author contributions

K.T. and Q.Z. contributed equally to this work. K.T., W.H., and D.X. were responsible for the study design. W.H. developed the original theoretical model, K.T. derived the extended model for somatic SV detection and implement the associated algorithms. Q.Z. reviewed the code and performed the evaluation of the algorithms. Y.L. and J.T. designed the experiments, J.W. executed the experiments. Q.Z. , L.F.Y. and Y.C.L. performed the simulation and visualization of data. L.X. analyzed the application scenarios of the algorithms and contributed to the discussion of their implications. All authors reviewed and approved the final manuscript.

## Competing interests

The authors declare no competing interests.

## Supplementary Figures

**Supplementary Figure S1.**
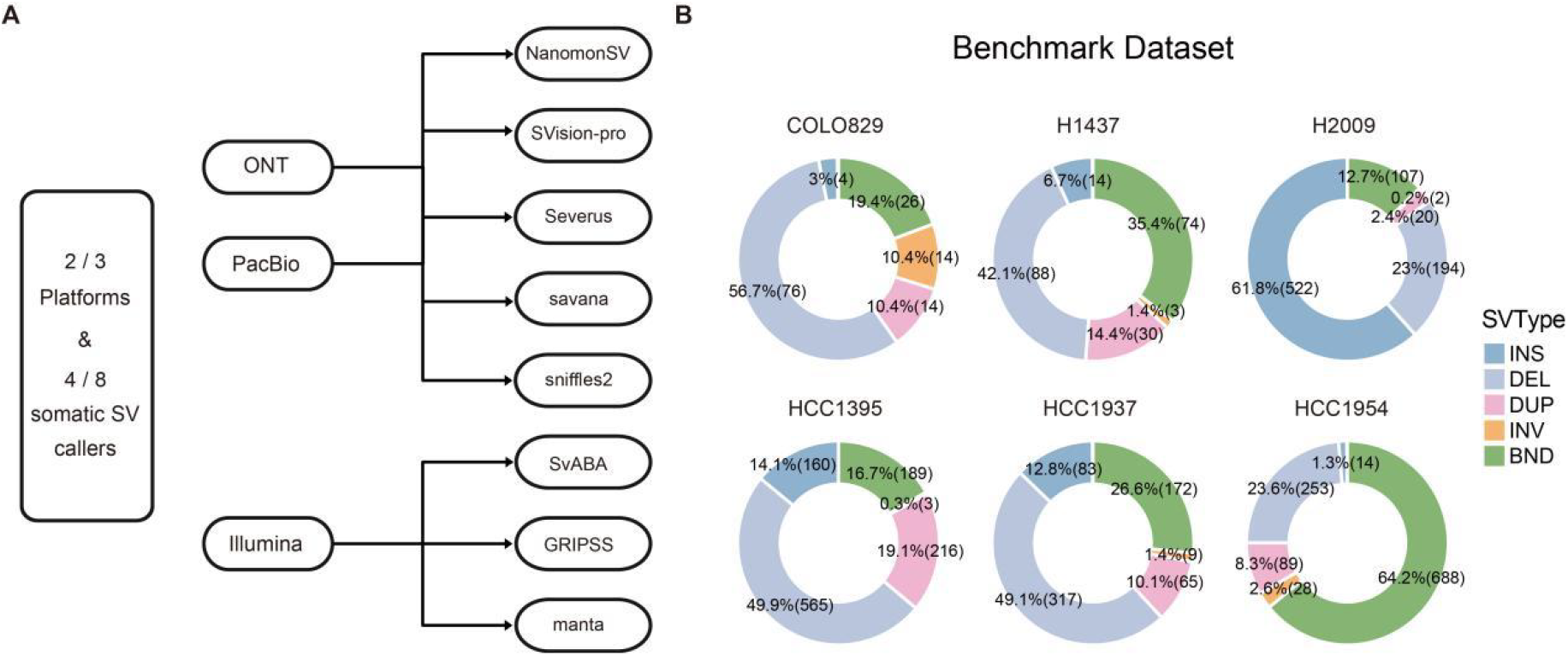
Construction workflow and structural variation distribution of benchmark datasets. A.Voting flowchart for benchmark dataset generation, illustrating the integration and filtering process of multi-method results. B. Pie charts depicting the distribution of somatic SV types across six cell line benchmark datasets. Data for H1437, HCC1395, HCC1937, H2009, and HCC1954 were generated through multi-omics voting consensus, while COLO829 data were derived from experimentally validated results in published literature (see Methods).

**Supplementary Figure S2.**
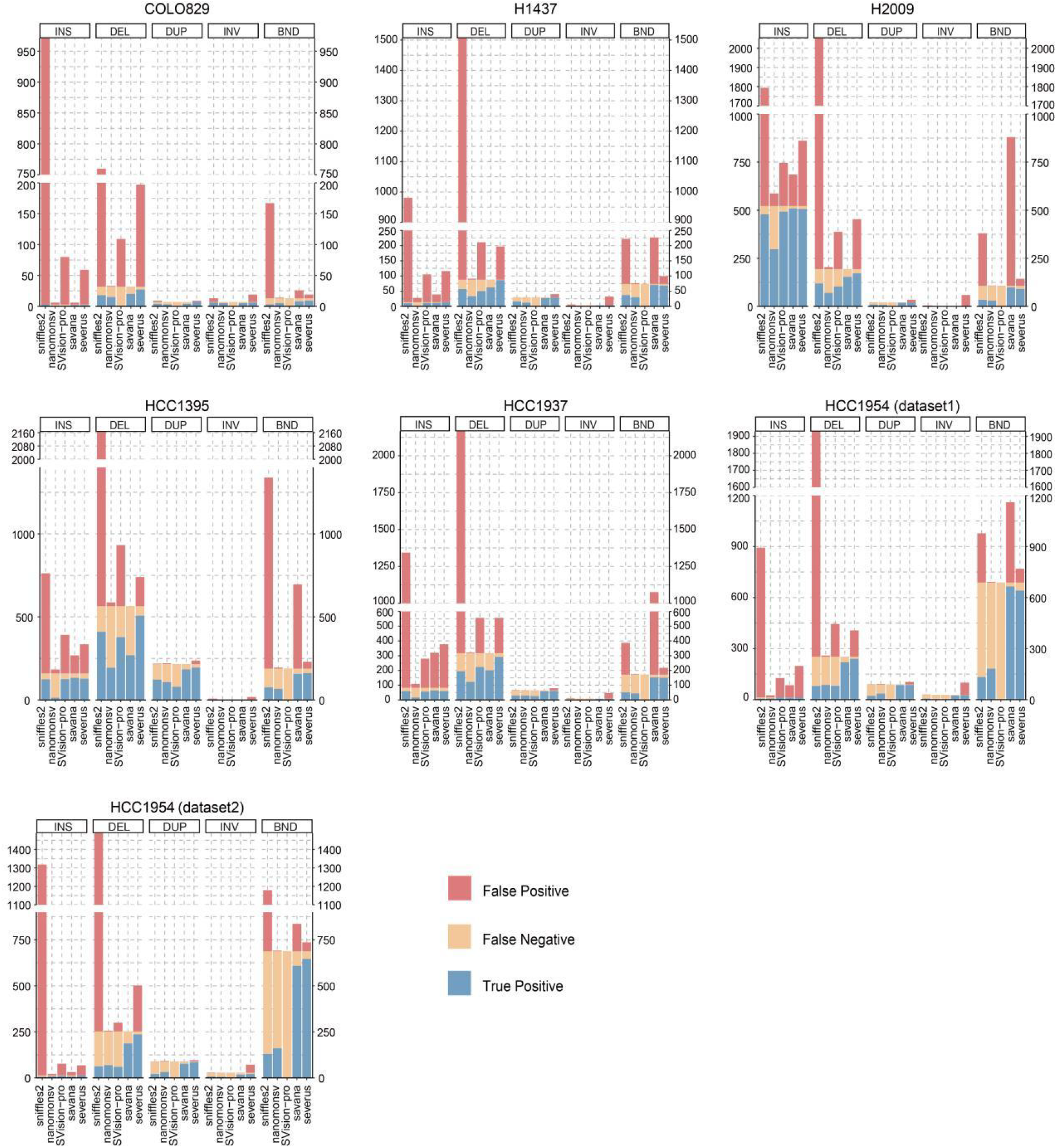
Performance comparison of five SV callers across six cell lines. Stacked bar charts showing the distribution of true positives (TP, blue), false positives (FP, red), and false negatives (FN, yellow) for each method. The X-axis represents the five methods (Sniffles2, nanomonsv, SVision-pro, savana, severus), and the Y-axis indicates the number of structural variations (SVs). Performance metrics are calculated relative to the benchmark datasets.

**Supplementary Figure S3.**
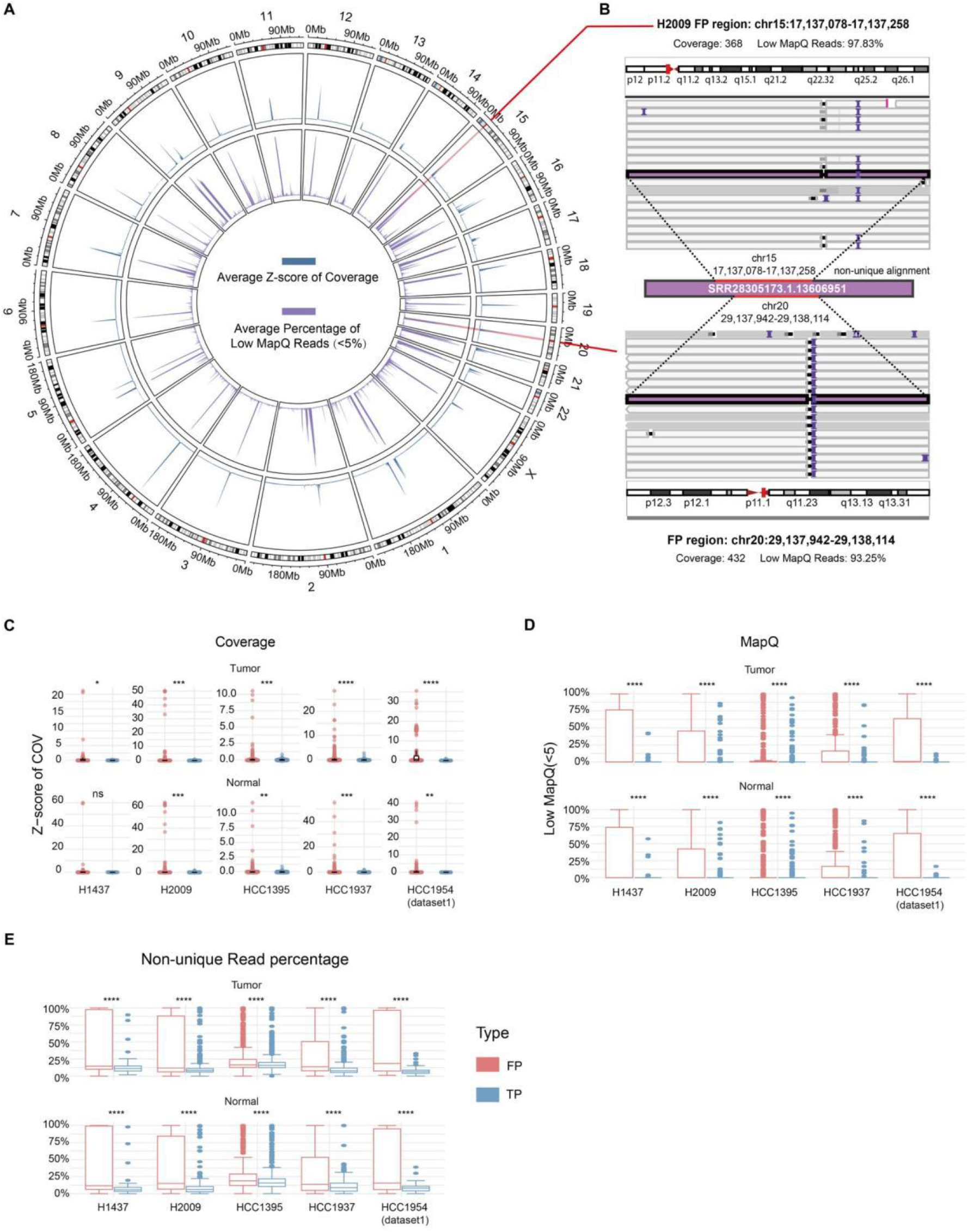
Genome-wide coverage, mapping quality and alignment ambiguity performance. A. Circle plot of chromosomes 1-22 and X displays three concentric layers: the outermost chromosomal karyotype bands, the middle blue line plot showing mean coverage Z-scores, and the innermost purple line plot indicating the mean proportion of low mapping quality (MAPQ <5) reads. B. IGV visualization of a non-unique alignment in H2009 cell line demonstrates a read simultaneously mapped to chr15:17137078-17137258 and chr20:29137942-29138114. C. Swarm plots compare coverage Z-scores between TP (blue) and shared FP (red) across five SV detection methods (Sniffles2, nanomonsv, SVision-pro, savana, severus) in tumor-normal cell line pairs, with statistical significance (t-test) annotated as ns (p > 0.05), * (p < 0.05), ** (p < 0.005), and *** (p < 0.0005). D-E. Box plots compare MAPQ values (D) and non-unique read percentages (E) between TP (blue) and shared FP (red).

**Supplementary Figure S4.**
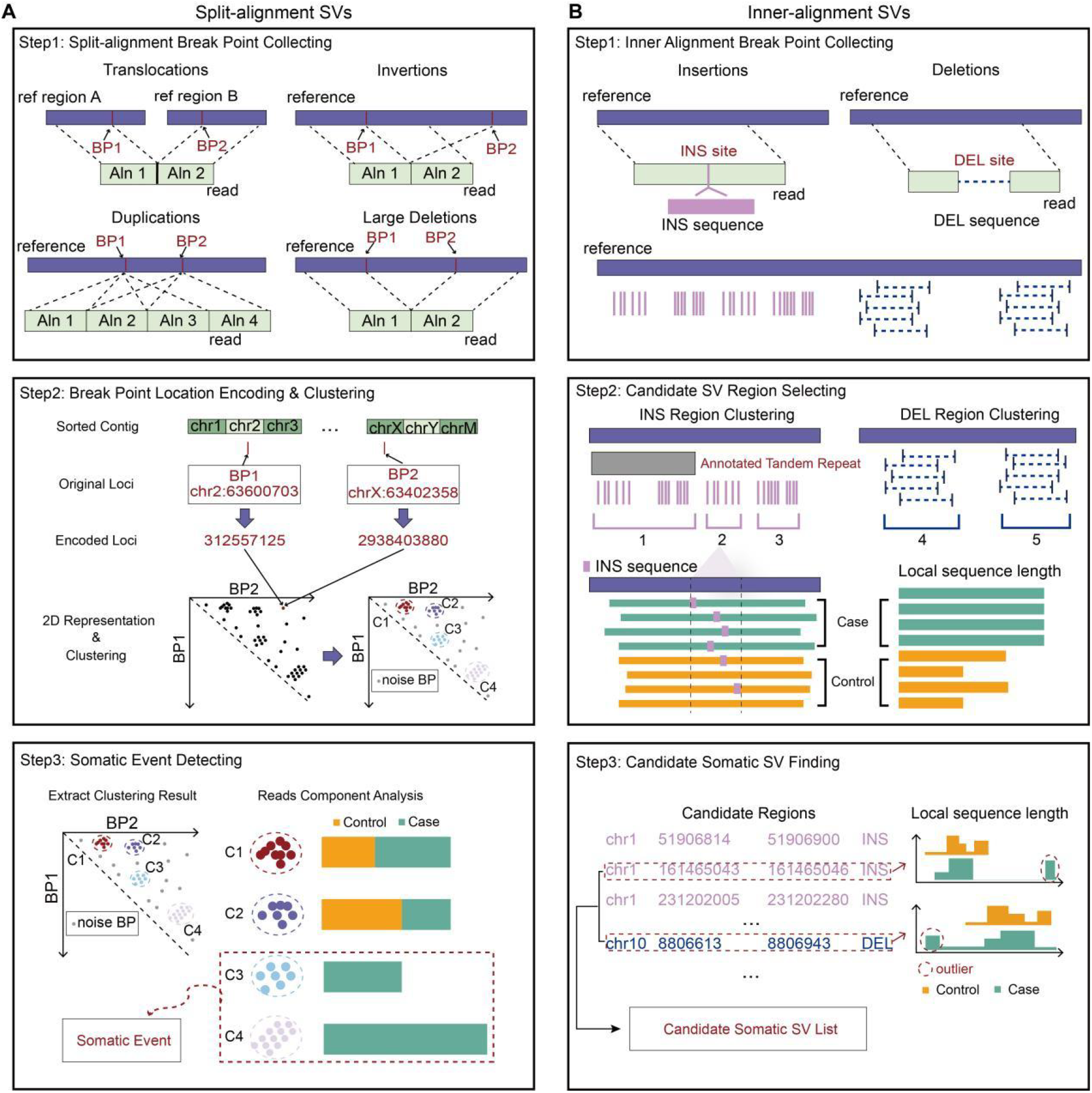
Detail Schematic of the candidate somatic SV selection workflow. A. Split-alignment SVs are selected based on paired-breakpoint DBScan clustering methods. B. Inner-alignment SVs are divided into insertion and deletions. Breakpoints of insertions are merged located within the same tandem repeat regions or within the length of distance threshold (by default 200bp). Breakpoints of deletions are directly merged if containing overlap. Full-length alignment information of each read from case and control samples are extracted to filter candidate inner-alignment SVs. Intervals with outlier reads more than 3 would be considered as candidate somatic intervals.

**Supplementary Figure S5.**
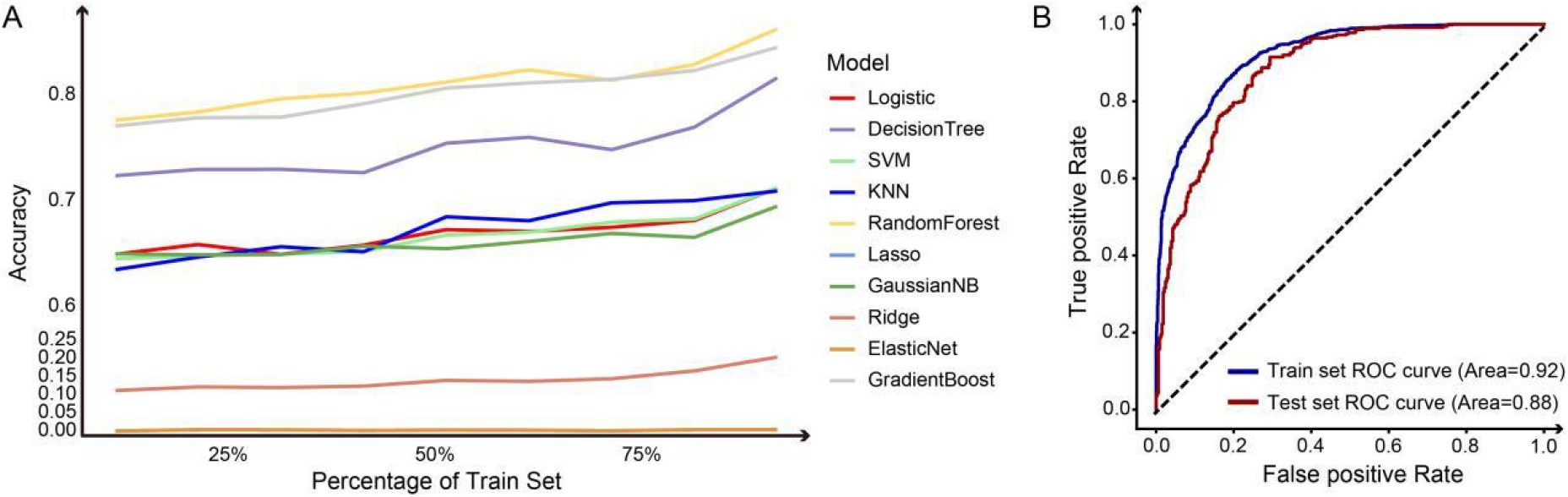
Machine learning model performance evaluation. A. Accuracy trend lines of ten machine learning methods: Logistic Regression, Decision Tree, Support Vector Machine (SVM), k-Nearest Neighbors (KNN), Random Forest, Lasso Regression, Gaussian Naive Bayes (GaussianNB), Ridge Regression, ElasticNet, and Gradient Boosting. Each method is represented by a distinct color. B. ROC curves of the Random Forest model. Blue curve: training set performance; red curve: test set performance.

**Supplementary Figure S6.**
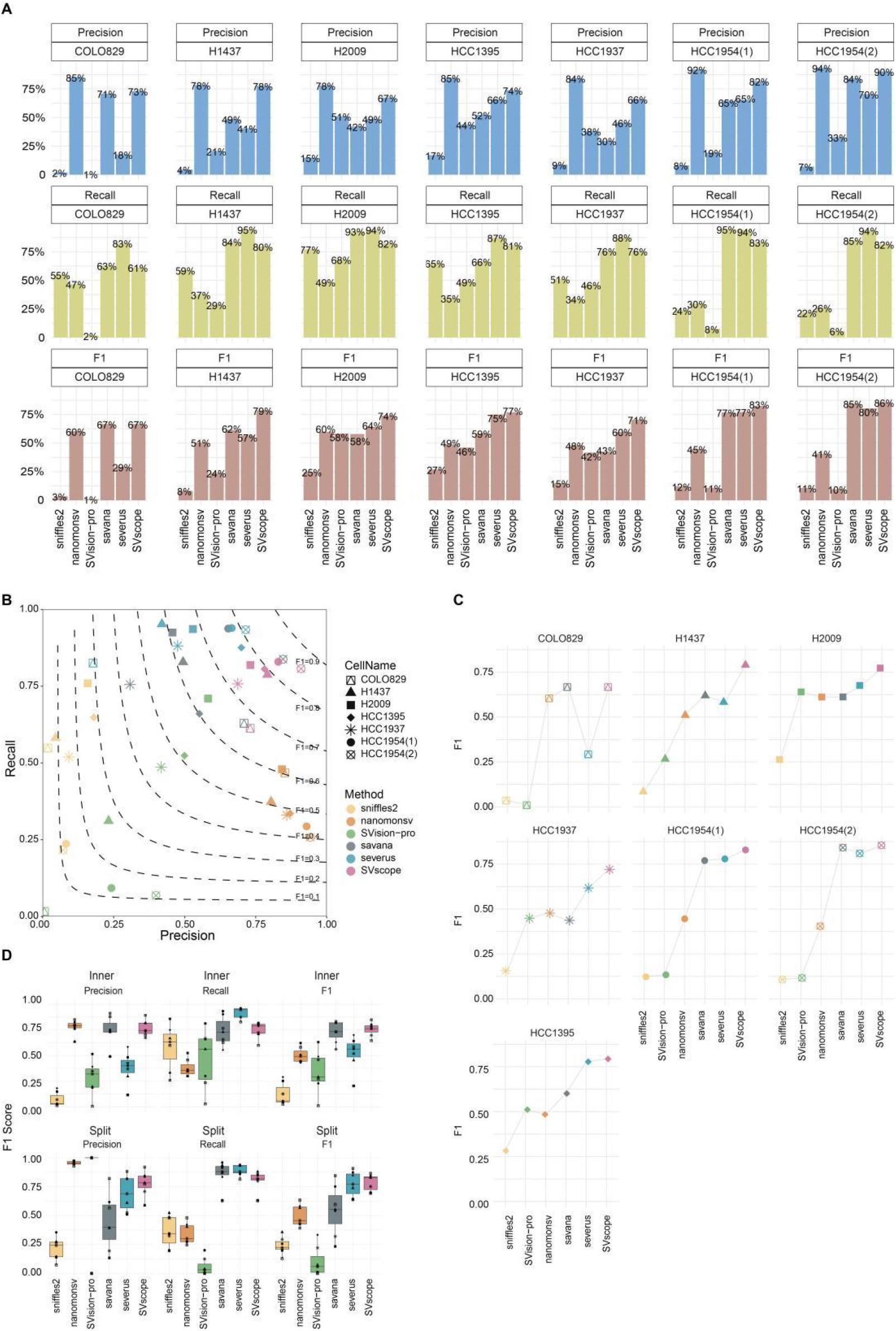
Performance comparison of SVscope and five other methods across seven cell lines. A. Stacked bar charts comparing Precision (blue), Recall (light green), and F1 score (brown) for SVscope and five other methods. X-axis: methods; Y-axis: percentage score. B. Scatter plot of Precision (X-axis) vs. Recall (Y-axis) across seven cell lines. Shapes denote cell lines; colors represent methods. Contour bands indicate F1 score levels (0.1 to 0.9). C. Line chart of F1 scores for SVscope and five methods across seven cell lines. X-axis: methods; Y-axis: F1 scores. Shapes correspond to cell lines; colors distinguish methods. D. Box plots comparing Precision, Recall, and F1 score for Inner SVs and Split SVs. X-axis: methods. Colors indicate methods; shapes mark cell lines.

**Supplementary Figure S7.**
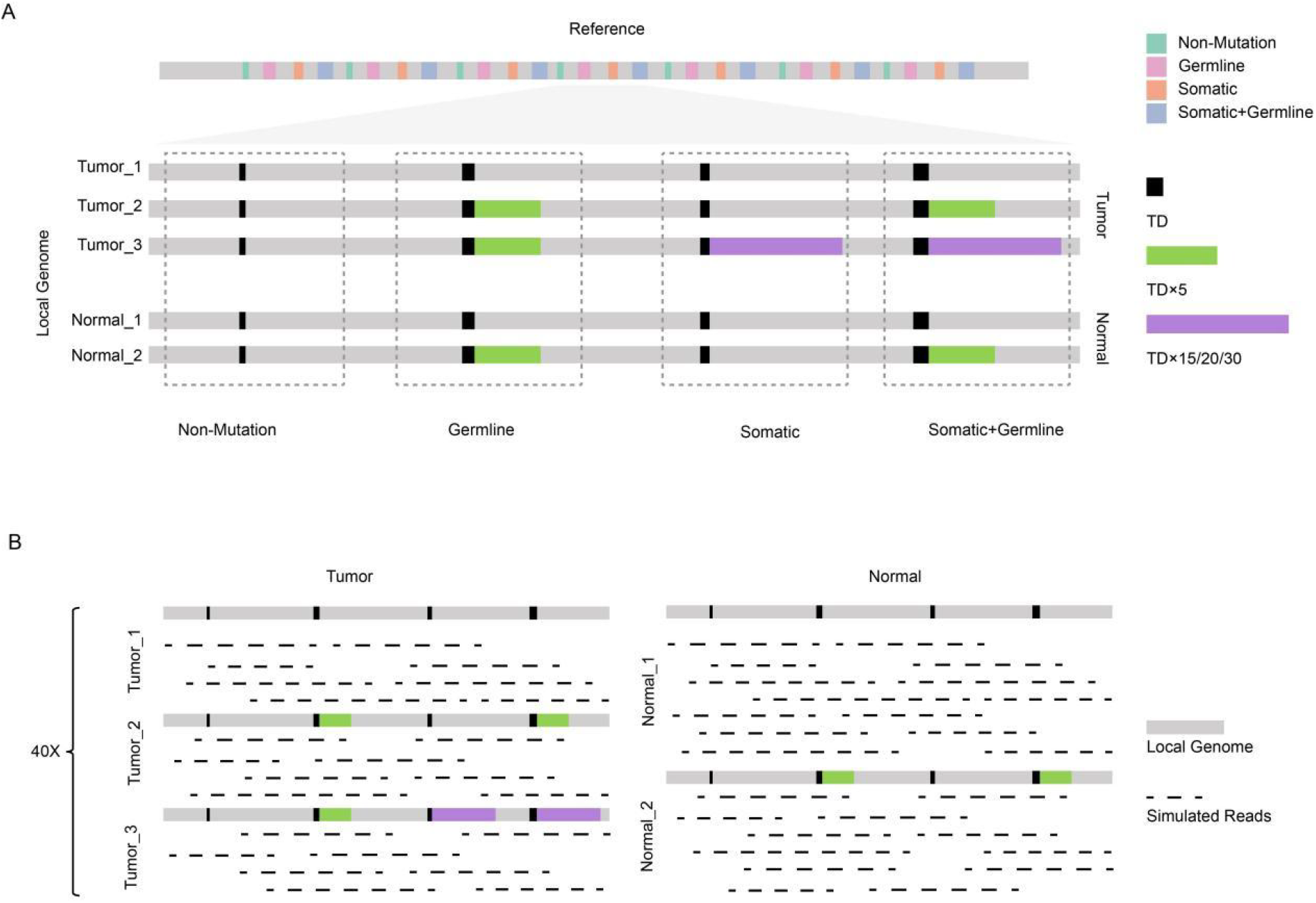
Schematic design of simulated tumor-normal datasets. A. Reference genome with four types of tandem duplications (TDs): Non-mutation (green), Germline (pink), Somatic (orange), and Mixed Somatic/Germline (lavender). Simulated tumor and normal local genomes were generated for three case types (X15, X20, X30), varying in TD length and complexity. B. Five tumor purity levels (20%, 30%, 40%, 50%, 100%) were simulated by adjusting the sequencing depth ratio of tumor-to-normal local genomes.

**Supplementary Figure S8.**
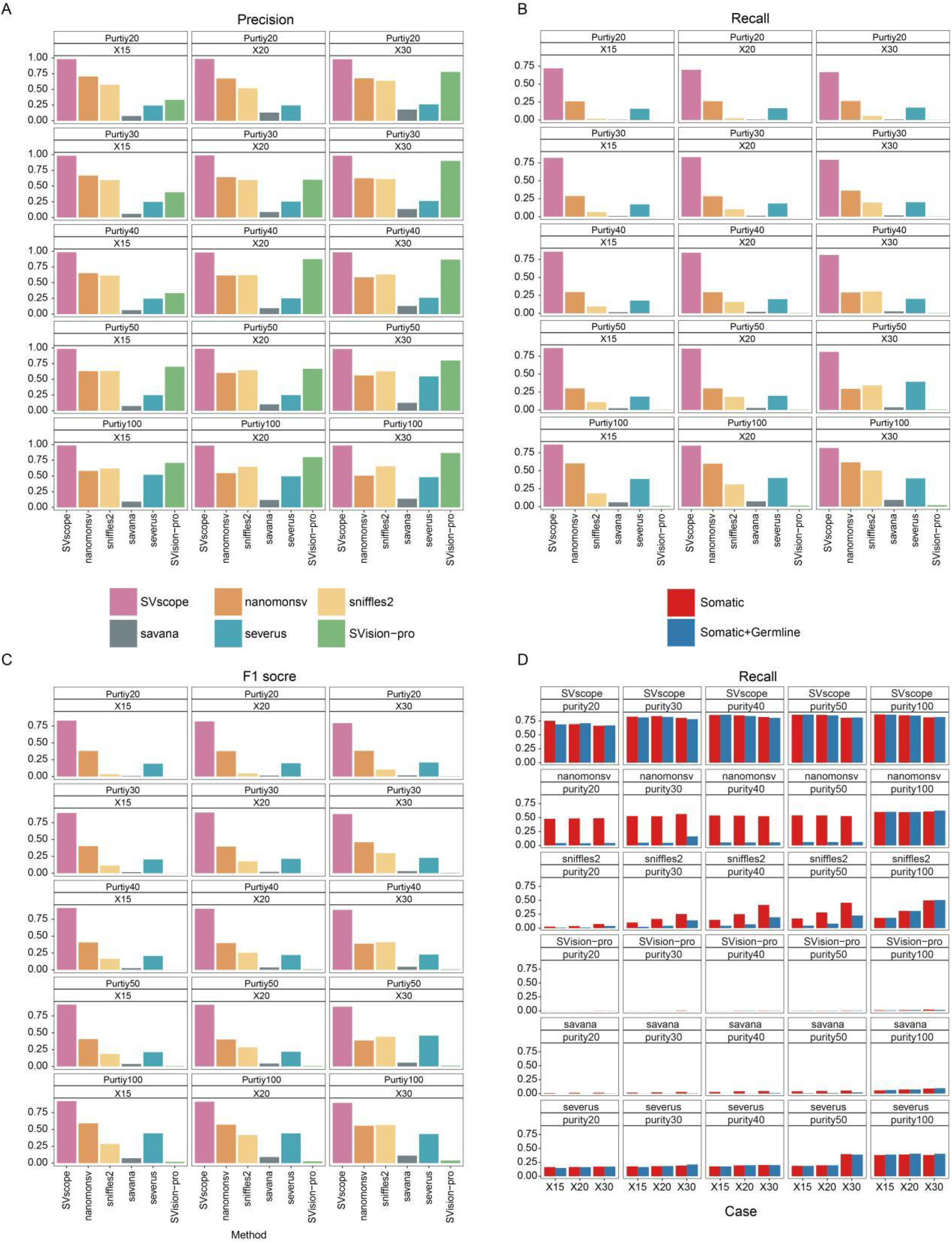
Performance evaluation of SV detection methods on simulated data. A-C. Bar plots compare Precision (A), Recall (B), and F1-score (C) across six methods: Sniffles2 (yellow), nanomonsv (orange), SVision-pro (green), savana (gray), severus (blue), and SVscope (pink). X-axis: methods; Y-axis: score percentages. Results are stratified by case (X15/X20/X30) and purity (20%, 30%, 40%, 50%, 100%). D. Recall comparison for Somatic (red) and Mixed Somatic/Germline (blue) across methods, cases, and purity levels. X-axis: methods; Y-axis: recall.

**Supplementary Figure S9.**
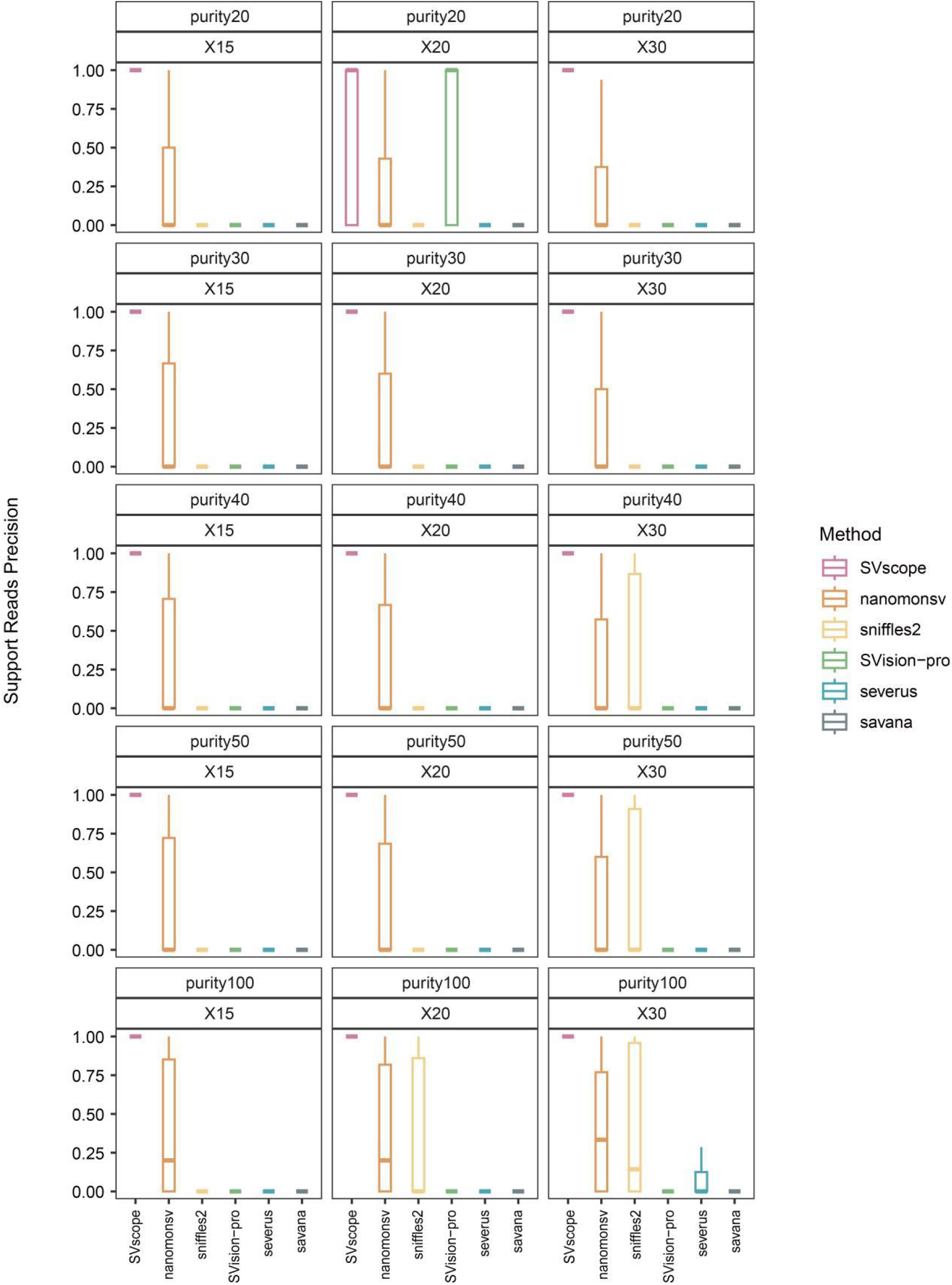
Support reads precision of SV detection methods on simulated data. Boxplots compare support reads precision (Y-axis: score) for six methods: Sniffles2 (yellow), nanomonsv (orange), SVision-pro (green), savana (gray), severus (blue), and SVscope (pink). X-axis labels represent methods, with box distributions aggregating results across three simulated cases (X15, X20, X30) and five tumor purity levels (20%, 30%, 40%, 50%, 100%).

**Supplementary Figure S10.**
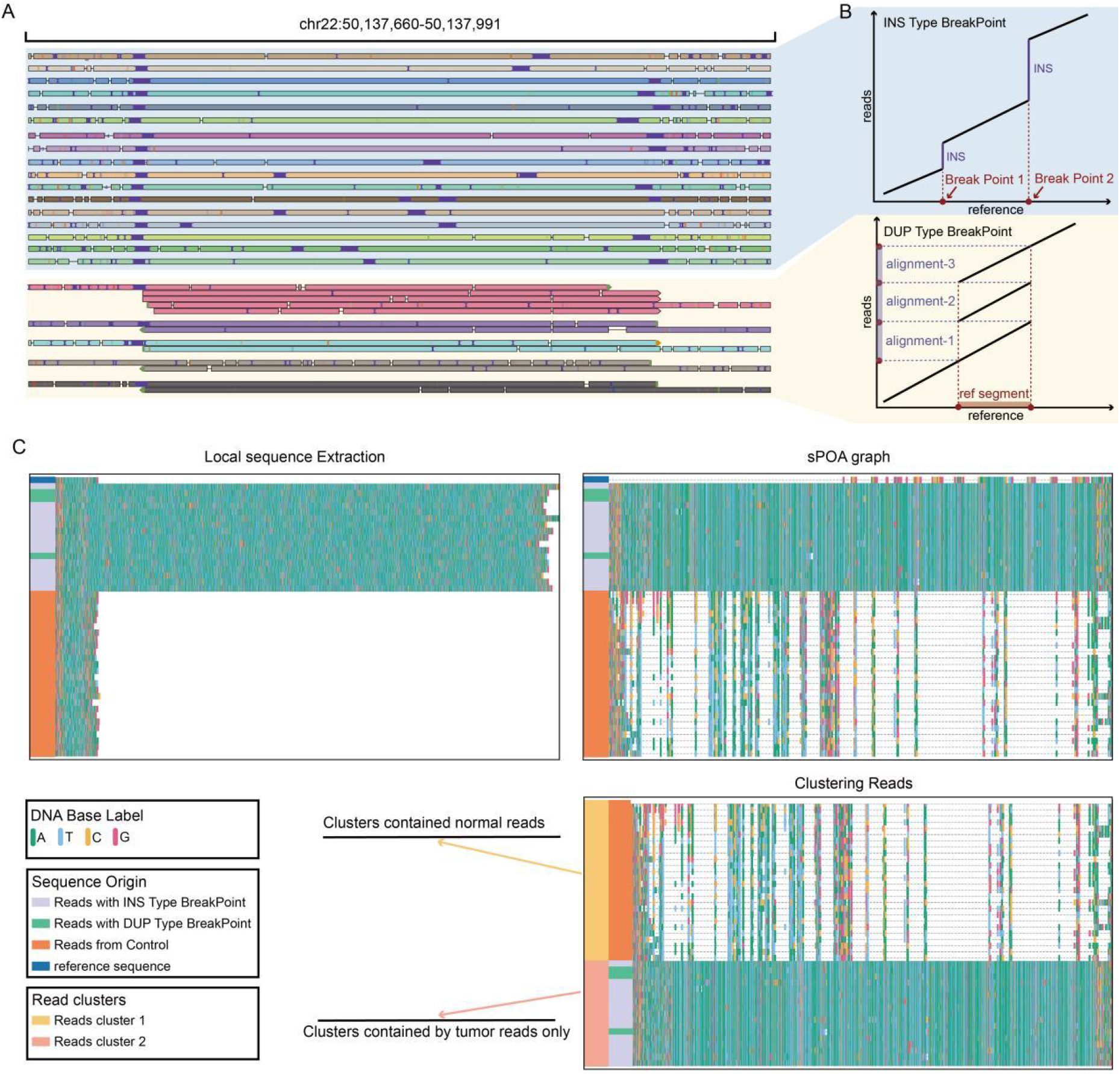
Representation of SVscope in heterogenous alignment breakpoint somatic SV case on chr22:50137660-50137991. A. IGV screen for simulated reads harboring somatic insertions in chr22:50137660-50137991. B. Line plot representing the alignment detail of INS type breakpoint (blue) and DUP type breakpoint (yellow). C. ScopeVIZ plot representing the process of local graph-genome optimization process of SVscope.

**Supplementary Figure S11.**
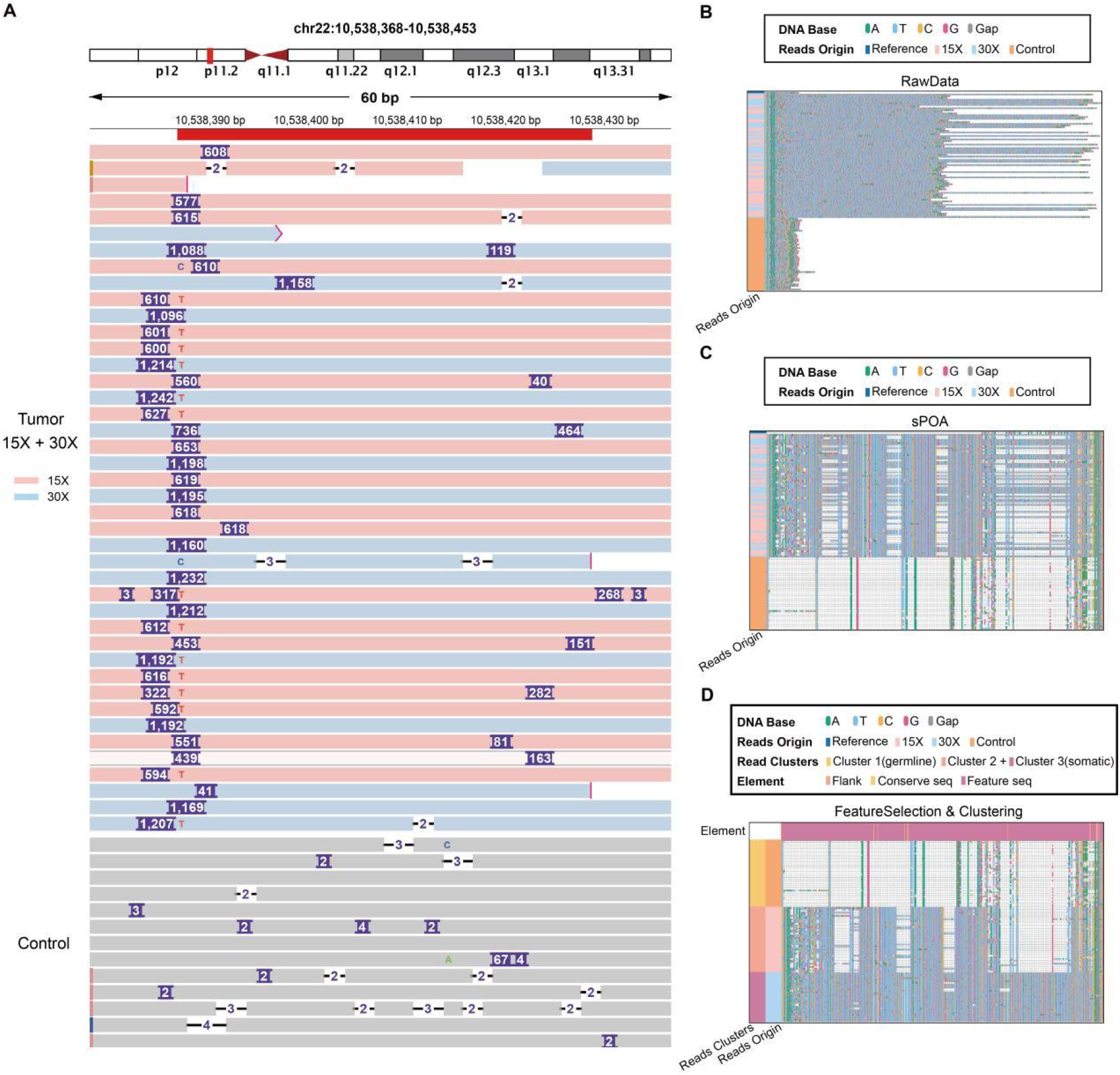
Integrated visualization and ScopeVIZ analysis of simulated structural variations in the chr22:10,538,368-10,538,453 region. A. IGV visualization of tumor (upper panel) and normal (lower panel) read alignments in the chr22:10,538,368-10,538,453 region. Tumor reads are a mixture of simulated X15 (pink) and X30 (blue) tandem duplications; normal reads (orange) lack these alterations. B. Sequence-level view of reads: Nucleotides are color-coded (A=green, T=cyan, C=yellow, G=magenta; gaps=gray dashes). Read origins are labeled (X15=pink, X30=blue, normal=orange). C. Partial order alignment graph generated by sparse partial order alignment (sPOA) from raw reads. D. SVscope-clustered reads: Top track classifies bases as Feature (magenta), Common (yellow), or Flank (peach). Left labels indicate SVscope-defined clusters(1/2/3) and read origins (X15/X30/normal).

**Supplementary Figure S12.**
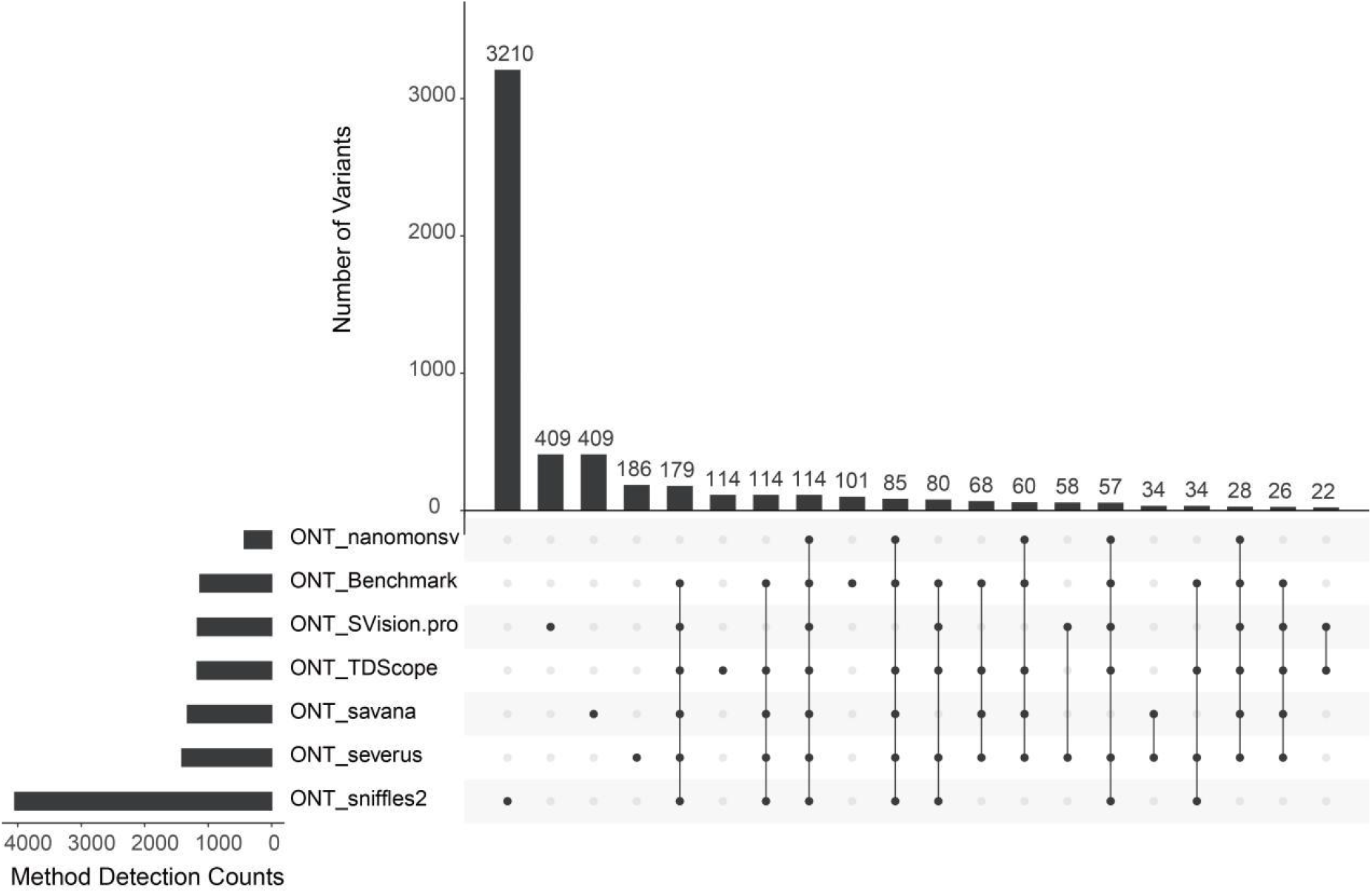
Overlap of structural variants (SVs) detected in the HCC1395 cell line. The upset plot displays intersections among SV calls from SVscope, five methods (Sniffles2, nanomonsv, SVision-pro, savanna, severus), and a benchmark dataset. The left bar plot shows the total SV counts per method and the benchmark dataset. The upper bar plot ranks the top 20 largest intersections by size (number of overlapping SVs). The central matrix uses dots to indicate method/dataset participation in each intersection (rows: methods/dataset; columns: intersection sets).

**Supplementary Figure S13.**
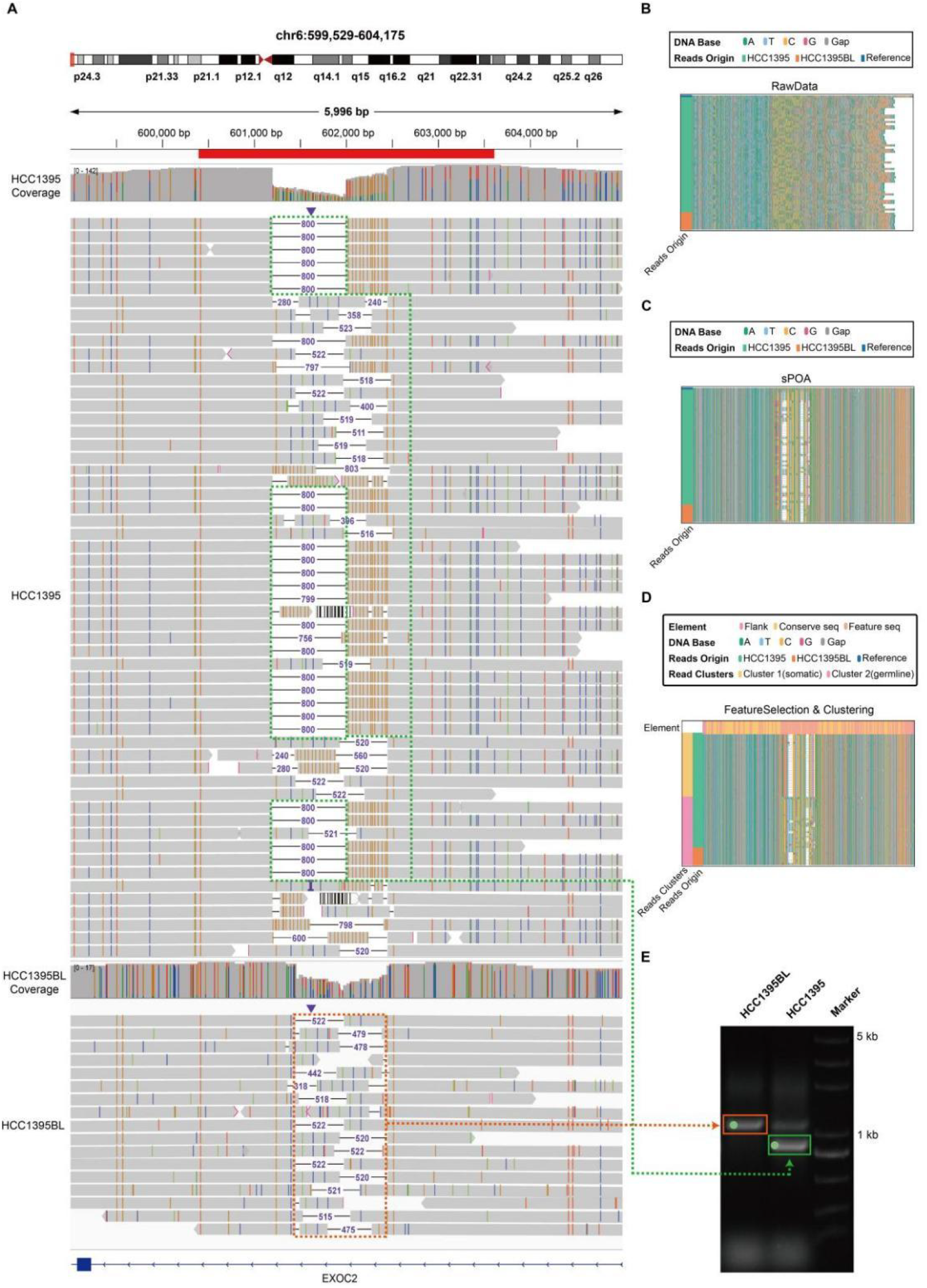
Detailed characterization of somatic SV and validation of somatic tandem repeat SV events of HCC1395 cell line in chr6:599,529-604,175. A. Genome browser plot shows the example somatic SV region chr6:599,529-604,175. in HCC1395 (top) and HCC1395BL(bottom). B. Bar plot shows the raw sequence extracted from HCC1395BL (colored in orange) and HCC1395 reads (colored in green). C. Bar plot shows multiple sequence alignment representation of sPOA graph based on read sequences from HCC1395BL (colored in orange) and HCC1395 reads (colored in green). D. SVscope-clustered reads: Top track classifies bases as Feature (magenta), Common (yellow), or Flank (peach). Left labels indicate SVscope-defined clusters(1/2) and read origins (tumor/normal). E. The gel electrophoresis image illustrates the targeted PCR results for the somatic SV event. Notably, the target bands are highlighted with green dots, clearly indicating the presence of the somatic SV event in both genomes.

**Supplementary Figure S14.**
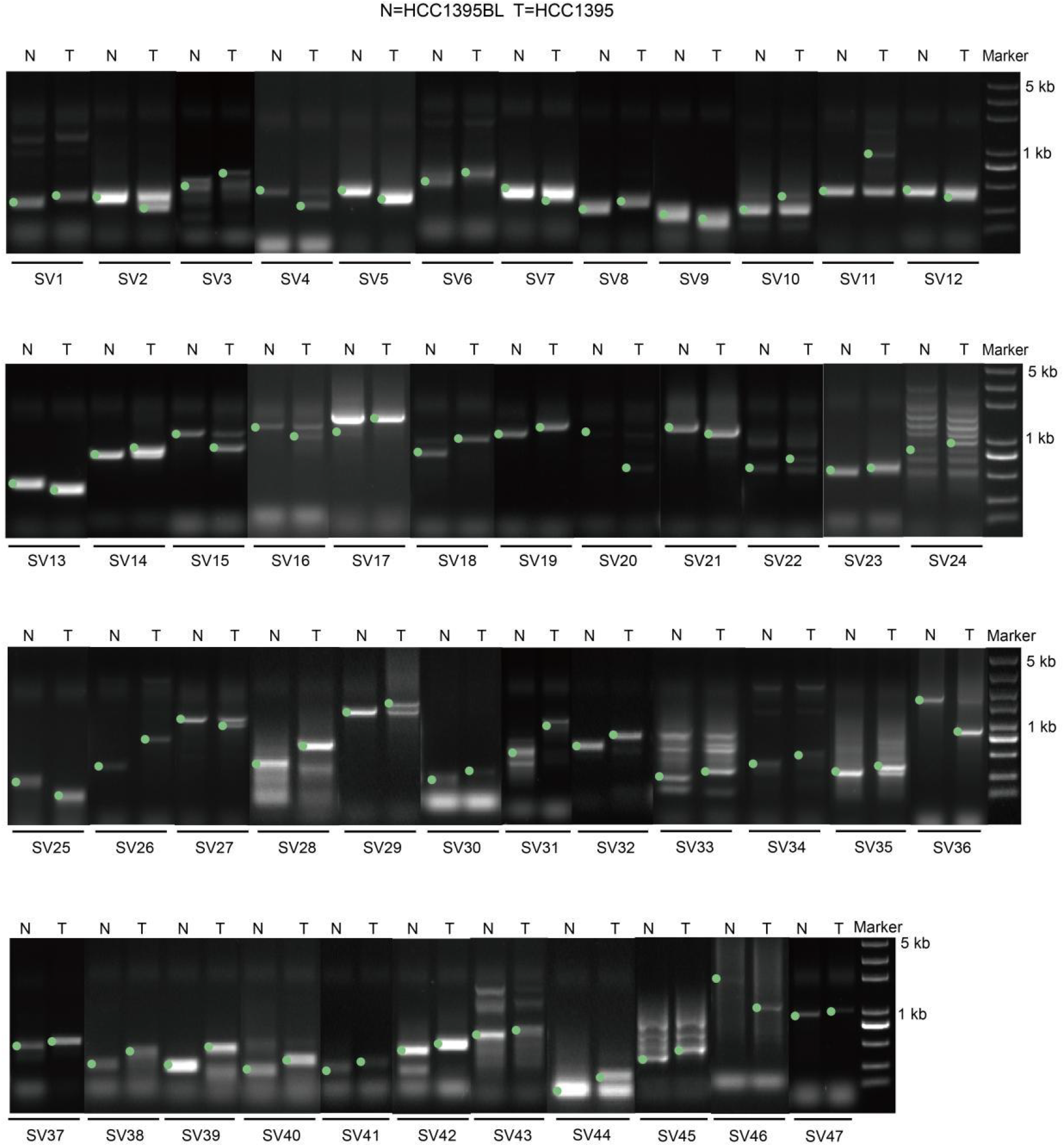
Gel electrophoresis validation of somatic structural variations (SVs) in the HCC1395 cell line. The gel electrophoresis image illustrates the targeted PCR results for 47 somatic SV events exclusively detected by the SVscope algorithm in the genomes of HCC1395BL (left panel, N) and HCC1395 (right panel, T). Notably, the target bands are highlighted with green dots, clearly indicating the presence of somatic SVevents in both genomes.

## Notes

### Competing Interest Statement

The authors have declared no competing interest.

### Summary of Updates

Update the title of paper. Update the function of original somatic SV detector. Rename the somatic SV detector as SVScope. Supplemental files updated.

